# Benchmarking Tools for Identification of rRNA Modifications in *Escherichia coli* using Oxford Nanopore Direct RNA Sequencing

**DOI:** 10.64898/2026.04.15.718756

**Authors:** Bhargava Reddy Morampalli, Olin K. Silander

## Abstract

RNA modifications are important for RNA structure, stability, and ribosome function, but their identification and localisation remains challenging. Oxford Nanopore direct RNA sequencing (DRS) enables modification-agnostic detection in native RNA, but existing tool benchmarks have focused almost exclusively on m6A in eukaryotic mRNA, leaving multi-modification tool performance in bacterial systems largely untested. Here, we benchmark ten RNA modification detection tools spanning signal-comparison, error-rate, and hybrid approaches on *Escherichia coli* K-12 MG1655 16S and 23S rRNA, which harbour 11 and 25 known modified sites, respectively, across 17 modification types. Using native RNA and *in vitro* transcribed (IVT) unmodified RNA, we evaluate performance across 25 coverage levels (5× to 1000×). DiffErr and JACUSA2 showed the strongest discrimination performance (AUROC >0.9 on both 16S and 23S rRNA), with DiffErr achieving the highest F1 score on 16S and JACUSA2 showing the most consistent precision-recall balance across both rRNAs. Both tools achieved full transcript-wide scoring and, along with DRUMMER, exact positional localisation. Several other tools produced no output at many rRNA positions, and restricting evaluation to reported positions inflated apparent performance. Signal-based tools showed a systematic 1-4 nucleotide 5ʹ offset from known modified positions, consistent with the ∼5-mer nucleotide stretch present in the read head of the nanopore; applying tool-specific offset corrections substantially improved per-site recovery and reduced false positives, substantially improving the performance of tools such as EpiNano and nanoDoc. At single-site resolution, no known modified site was recovered by all tools, and several m5C, m5U, and m6A sites were missed by the majority of tools. Tool combination analysis showed that pairing error-rate-based tools with offset-corrected signal-based tools improved site recovery beyond any individual tool, with the best three-tool combination recovering 30 of the 36 known sites while maintaining low false positive rates. These results establish that discrimination metrics (e.g. AUROC) alone are insufficient to evaluate modification detection tools: output completeness, positional precision, and per-modification-type sensitivity should be reported alongside standard benchmarking metrics.

## Introduction

Covalent RNA modifications have been implicated in a number of important cellular processes in bacteria, including translation rate and termination (Hoernes et al., 2016), translation accuracy (Hoernes et al., 2016), RNA folding and stability (Durant et al., 2005), responses to antibiotic stress (Babosan et al., 2022) and environmental stress (Fleming et al., 2023b). N6-methyladenosine (m^6^A) and pseudouridine (Ψ) are among the most abundant and well-studied modifications, though the largest number of modifications occur in transfer RNAs (tRNA) and ribosomal RNAs (rRNA) (Sordyl et al., 2026). More than 170 different modifications have been identified across all kingdoms of life (Xuan et al., 2024; Sordyl et al., 2026). Identifying the location, type, and frequency of different modifications not only defines the modification landscape but may also yield mechanistic insights into how they mediate phenotypic changes.

Traditional methods of identifying RNA modifications include liquid chromatography-mass spectrometry (LC-MS) (Antoine et al., 2019), antibody-based immunoprecipitation (Dominissini et al., 2016) and chemical or enzymatic conversion protocols targeting individual modifications such as bisulfite sequencing for m^5^C (Schaefer et al., 2009), AlkAniline-Seq for m^7^G and m^3^C (Marchand et al., 2018), miCLIP-seq for m^6^A modifications (Grozhik et al., 2017), and N_3_-CMC-based method for pseudouridine (X. Li et al., 2015). Most of these methods are restricted to one type of modification per experiment and several lack single-nucleotide resolution, which makes them unsuitable for studying multiple modification types simultaneously across the transcriptome.

In contrast to other sequencing technologies, the Oxford Nanopore Technologies (ONT) sequencing platform is capable of direct RNA sequencing (DRS). As with ONT DNA sequencing, the change in ionic current during the translocation of RNA through a pore depends on shape and charge of the molecule being translocated, leading to differences in signal for modified and unmodified ribonucleotides (Anreiter et al., 2021; Jain et al., 2022). Because the nanopore read head interrogates a window of approximately five nucleotides at any given time (White & Hesselberth, 2022), modifications can cause signal differences in the surrounding region of the exact modified site, which can lead to misidentified positions if this is not accounted for. DRS has been applied to several bacterial systems, such as transcriptome characterisation in *E. coli* (Fleming et al., 2023a; Guo et al., 2025; Riquelme-Barrios et al., 2025) and Pseudomonas aeruginosa (Pust et al., 2022). In *E. coli,* 16S and 23S rRNAs together harbour 36 known modification sites spanning 17 distinct chemistries (Popova & Williamson, 2014; Fleming et al., 2023a), making them one of the most comprehensively annotated modification landscapes in any organism and an ideal substrate for multi-modification type benchmarking.

Several tools have been developed to identify RNA modifications from ONT DRS data, and their number has grown rapidly, from approximately 15 tools reviewed in 2022 (White & Hesselberth, 2022; Zhao et al., 2022) to over 80 evaluated (original tools and retrained versions) by 2026 (Luo et al., 2026). In general, they use one of three methods to identify modifications: contextualising basecalling errors to identify modifications (Jenjaroenpun et al., 2021; Liu et al., 2019; Naarmann-de Vries et al., 2023; Parker et al., 2020; Abebe et al., 2022); the development of new basecalling models to identify modified nucleotides (Cruciani et al., 2023; Diensthuber et al., 2023); and direct comparison between the signals of modified and unmodified RNA molecules (Begik et al., 2021; Gao et al., 2021; Hassan et al., 2021; Leger et al., 2021; Lorenz et al., 2020; Lucas et al., 2024; Maier et al., 2020; Parker et al., 2021a; Pratanwanich et al., 2021; Stoiber et al., 2017; Ueda, 2021). Some consensus methods rely on more than one of these approaches (Delgado-Tejedor et al., 2024). Recent work has further expanded the methodological landscape, including approaches for m6A detection and quantification at single-molecule resolution (Qin et al., 2022; Cruciani et al., 2025), improved basecalling algorithms and alternative model parametrisation (Diensthuber et al., 2023; Fonzino et al., 2024; Teng et al., 2024), and the application of deep learning to detect diverse modifications (Nguyen et al., 2022a; Mateos et al., 2024; Martinek et al., 2024; Wu et al., 2024).

Error-based methods, while computationally efficient, are inherently prone to changes in accuracy because the accuracy of ONT basecalling itself can change. Modification-aware basecalling models require modification types and locations to be known in advance and are currently trained on a limited subset of modification chemistries, restricting their applicability to novel or diverse modification types. Tools based on direct comparison of raw nanopore signals between matched modified and unmodified control samples can achieve high accuracy for multiple modification types when appropriate controls are available, although they generally cannot identify the type of modification. Because they compare matched samples sequenced under the same conditions, signal-comparison approaches can in principle be more robust to global shifts in signal properties than models that assume a fixed canonical signal distribution, though systematic assessments across chemistries remain limited (Luo et al., 2026). However, signal comparison methods require a means to obtain unmodified RNA for comparison, through direct synthesis of DNA templates followed by i*n vitro* transcription (Nguyen et al., 2022b), gene cloning (Patiño-Guillén et al., 2024; Smith et al., 2020), PCR, or other enrichment (Leger et al., 2021) followed by *in vitro* transcription, or transcriptome-wide reverse transcription followed by *in vitro* transcription (Morampalli et al., 2021).

A number of recent publications have benchmarked RNA modification detection tools, including review of computational methods, supported modification types and reported performance metrics of 15 tools (Zhao et al., 2022), performance of ten tools on Sindbis virus (Tan, Guo, Wang, et al., 2024a), identification of m6A on *E. coli* and *S. aureus* mRNA (Tan, Guo, Shao, et al., 2024), evaluation of m6A detection accuracy (Zhong et al., 2023a), a comprehensive epitranscriptomic study of E. coli using DRS (Riquelme-Barrios et al., 2025), and a systematic evaluation of 86 tools across six modification types (Luo et al., 2026). However, these benchmarks have predominantly evaluated m6A detection in eukaryotic mRNA from human or mouse datasets (Zhong et al., 2023a; Maestri et al., 2024; Luo et al., 2026), m6A in bacterial mRNA (Tan, Guo, Shao, et al., 2024) or viral RNA (Tan, Guo, Wang, et al., 2024b), and the performance of current tools on diverse modification types in bacteria remains largely uncharacterised. Beyond these taxonomic and modification-type gaps, no study has systematically characterised tool output completeness (whether a score is produced for every nucleotide position in the molecule), positional precision, or per-modification-type sensitivity across a common set of tools and ground-truth sites. Finally, although sequencing depth strongly influences precision and recall (Maestri et al., 2024), the minimum coverage required for reliable detection and how coverage-dependent performance varies across tools and modification types has not been systematically quantified across tools.

Here we selected ten computational tools that require matched unmodified controls, an approach that generally maximises precision and sensitivity in modification detection (Cruciani & Novoa, 2025), requiring also that the tools have the potential to identify multiple modification types (Table 1). We evaluated these tools for their capacity to identify 36 known modification sites across 17 modification types on 16S and 23S ribosomal RNA (rRNA) of *Escherichia coli* K-12 MG1655 (**Supp. Tables S1-S2**) (11 sites on 16S rRNA and 25 sites on 23S rRNA).

**Table 1.**
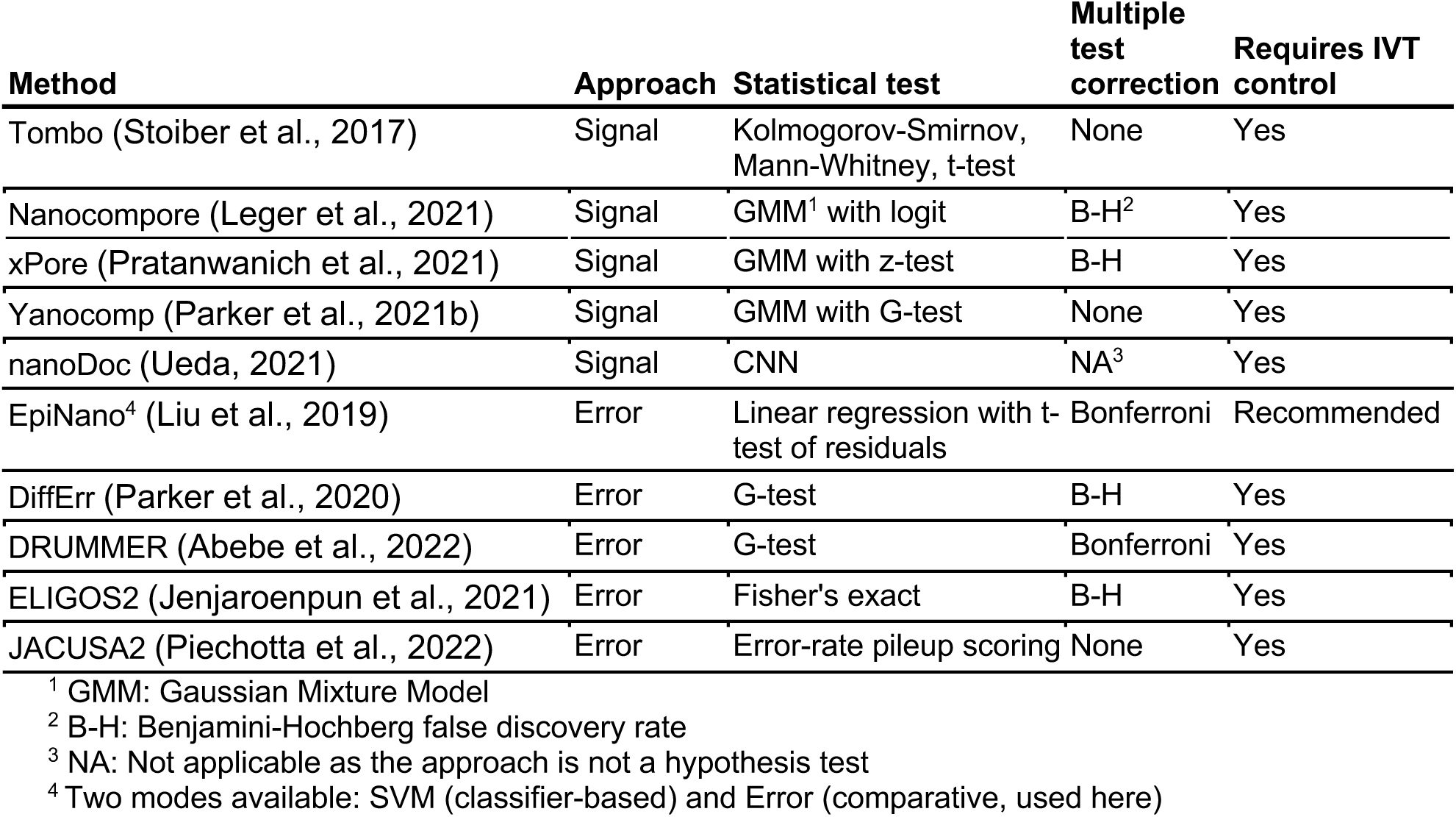
RNA modification detection tools benchmarked in this study.

Beyond standard discrimination metrics (AUROC, AUPRC), we characterised tool output completeness (fraction of positions scored by each tool), positional precision, number of false positives reported by each tool, and site-level detection patterns. We quantified these across 25 DRS coverage levels to determine how sensitivity, precision, and positional precision change as coverage decreases for each tool. We also test for systematic positional offsets and the effects of correcting for these offsets.

## Results

### Direct RNA sequencing of native and IVT controls yields high-coverage rRNA datasets

We isolated total RNA from three biological replicates of mid-exponential phase *E. coli* K-12 MG1655 cultures (native RNA; **Methods**), and generated matched IVT controls (IVT RNA; ***Methods***). We sequenced all six samples using the direct RNA sequencing protocol from ONT (SQK-RNA002 kit chemistry), obtaining 310,025 (replicate 1), 496,554 (replicate 2), and 648,245 (replicate 3) reads for native RNA. For IVT RNA we obtained 111,968 (replicate 1), 92,783 (replicate 2) and 90,981 (replicate 3) reads. Approximately 85% of the DRS reads mapped to the 16S and 23S rRNA regions.

Native rRNA reads spanned the full length of both the 16S (1,542 bp) and the 23S (2,904 bp), although only a small fraction of 23S reads were full-length (**Fig. 1**). Read lengths were multimodal in both native and IVT datasets, with a substantial fraction of short reads (**Fig. 1**). IVT reads were consistently shorter than native reads, likely due to truncation during reverse transcription or T7 transcription, and we observed almost no full-length IVT transcripts. Read quality was positively correlated with read length for both native and IVT RNA datasets, and at equivalent read lengths, native and IVT quality scores were comparable (**Supp. Fig. S2**).

**Figure 1.**
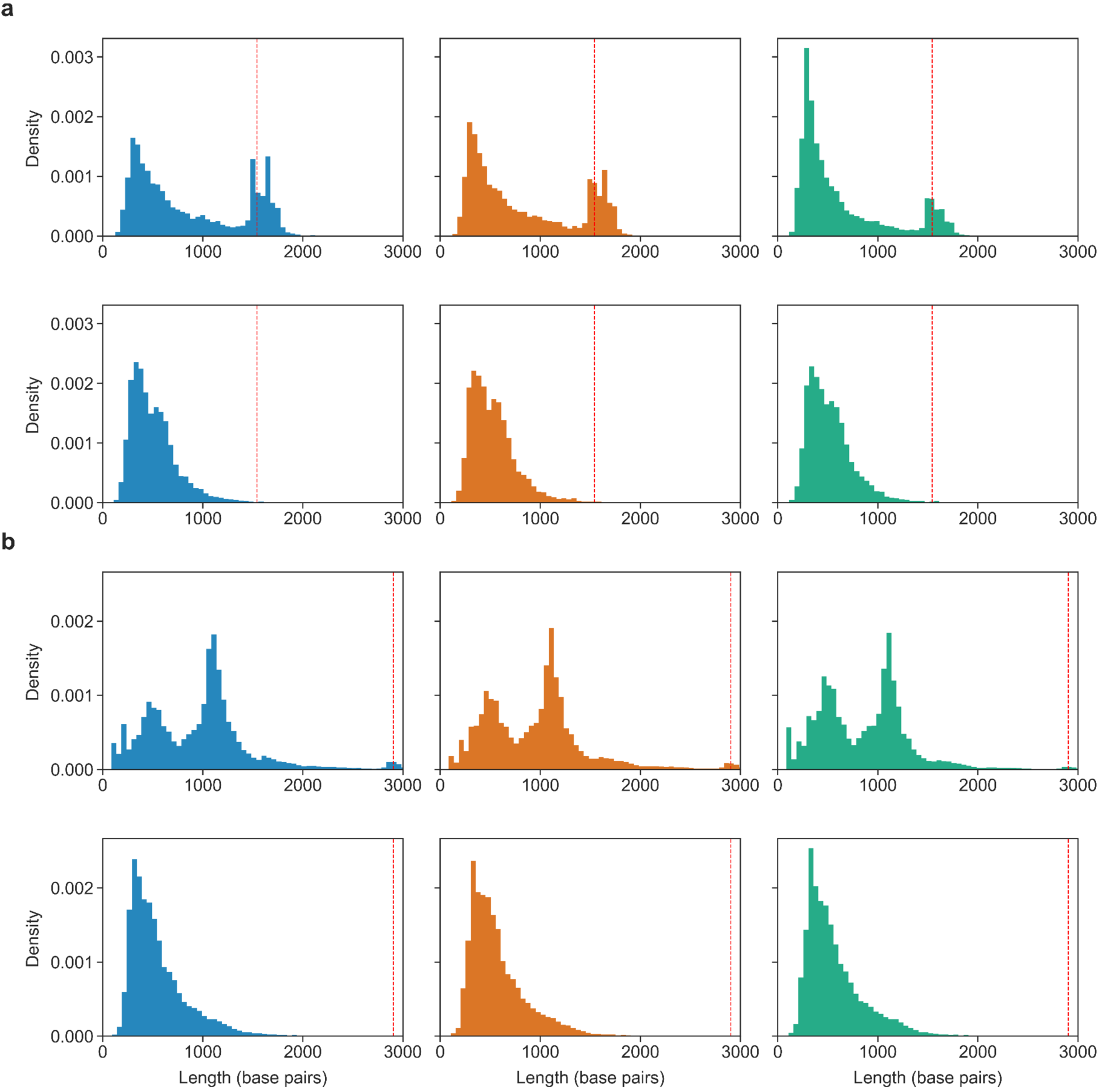
IVT rRNA reads are shorter than native rRNA reads. Read length distributions for native (top row) and IVT (bottom row) rRNA for (**a)** 16S and (**b)** 23S. Dashed red lines indicate the full-length transcript (1,542 bp for 16S; 2,904 bp for 23S). Colours indicate biological replicates (blue, orange, green). Native reads include a substantial fraction of full-length transcripts for 16S but not 23S; IVT reads are consistently shorter. rRNA is co-transcribed with tRNA and/or 5S rRNA making some transcripts longer than the canonical 16S or 23S rRNA length.

Coverage was strongly biased toward the 3ʹ end of both rRNA transcripts, consistent with previous observations (Fleming et al., 2023a; Jain et al., 2022; Workman et al., 2019), and with the fact that RNA is sequenced in the 3’ to 5’ direction. Coverage for native 16S rRNA was approximately three-fold higher at the 3ʹ end than at the 5ʹ end. For IVT 16S rRNA, this difference was up to 100-fold (**Fig. 2a**). For 23S rRNA, both native and IVT coverage differed by only 10-fold between the 5ʹ and 3ʹ ends (**Fig. 2b**). Despite the lower coverage of IVT RNA, the rRNA regions still achieved up to 35,000-fold coverage.

**Figure 2.**
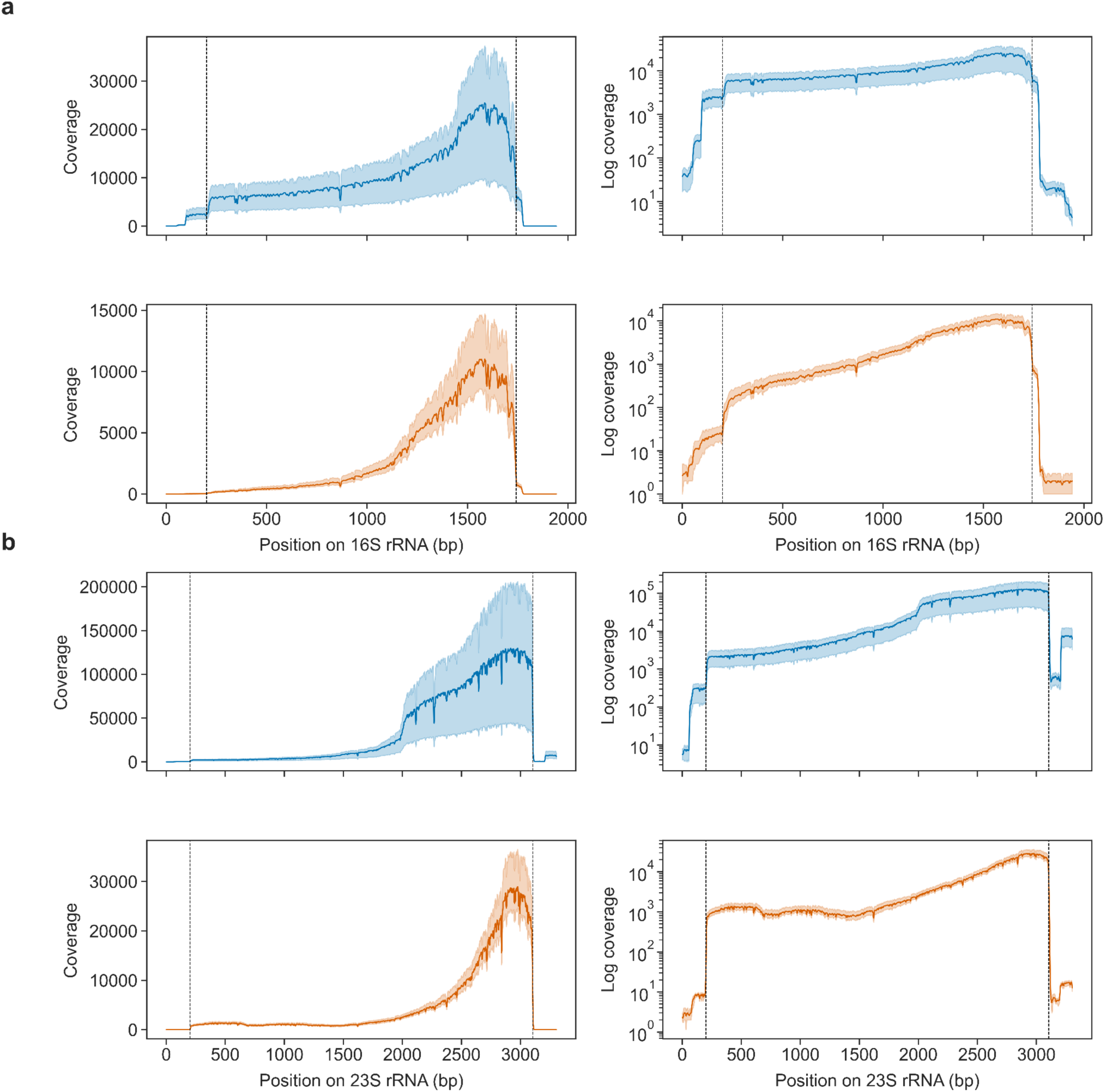
Direct RNA sequencing coverage is biassed toward the 3ʹ end. Coverage across (**a)** 16S and (**b)** 23S rRNA for native (blue, top row) and IVT (orange, bottom row) reads. Left panels use a linear y-axis; right panels use a log y-axis. Solid lines show mean coverage across three biological replicates; shaded bands indicate the replicate range (min-max). We smoothed coverage values over a window of 7 bp to mitigate artifactual dips from systematic basecalling indels. Dashed vertical lines mark the annotated 5’ and 3’ ends of each rRNA gene (positions 1 and 1542 for 16SrRNA; positions 1 and 2904 for 23S rRNA); 200 nt of flanking sequence on each side was included in the mapping reference. The coverage increase beyond the 3’ end of 23S reflects the downstream 5S rRNA transcript.

### Coverage affects the accuracy of modification detection

To investigate the effect of sequencing coverage on the accuracy of modification detection, we subsampled the native and IVT rRNA sequencing data to 25 different coverage levels ranging from 5× to 1000× (see **Methods**), (**Supp. Fig. S1**) and benchmarked ten comparative computational tools (**Table 1**) on known RNA modifications on *E. coli* 16S and 23S rRNA across all coverage levels.

Seven of the ten tools (JACUSA2, ELIGOS2, Tombo, xPore, DiffErr, Nanocompore, and EpiNano) produced outputs across all 25 subsampled coverage levels for both rRNA species (**Fig. 3**; **Supp. Fig. S7**). DRUMMER required at least 25× coverage of 16S to provide output on probable site modification, or 15× for 23S. Yanocomp required at least 35× (16S) or 20× (23S) coverage. We tested nanoDoc at only 15 of the 25 coverage levels (see **Methods**).

**Figure 3.**
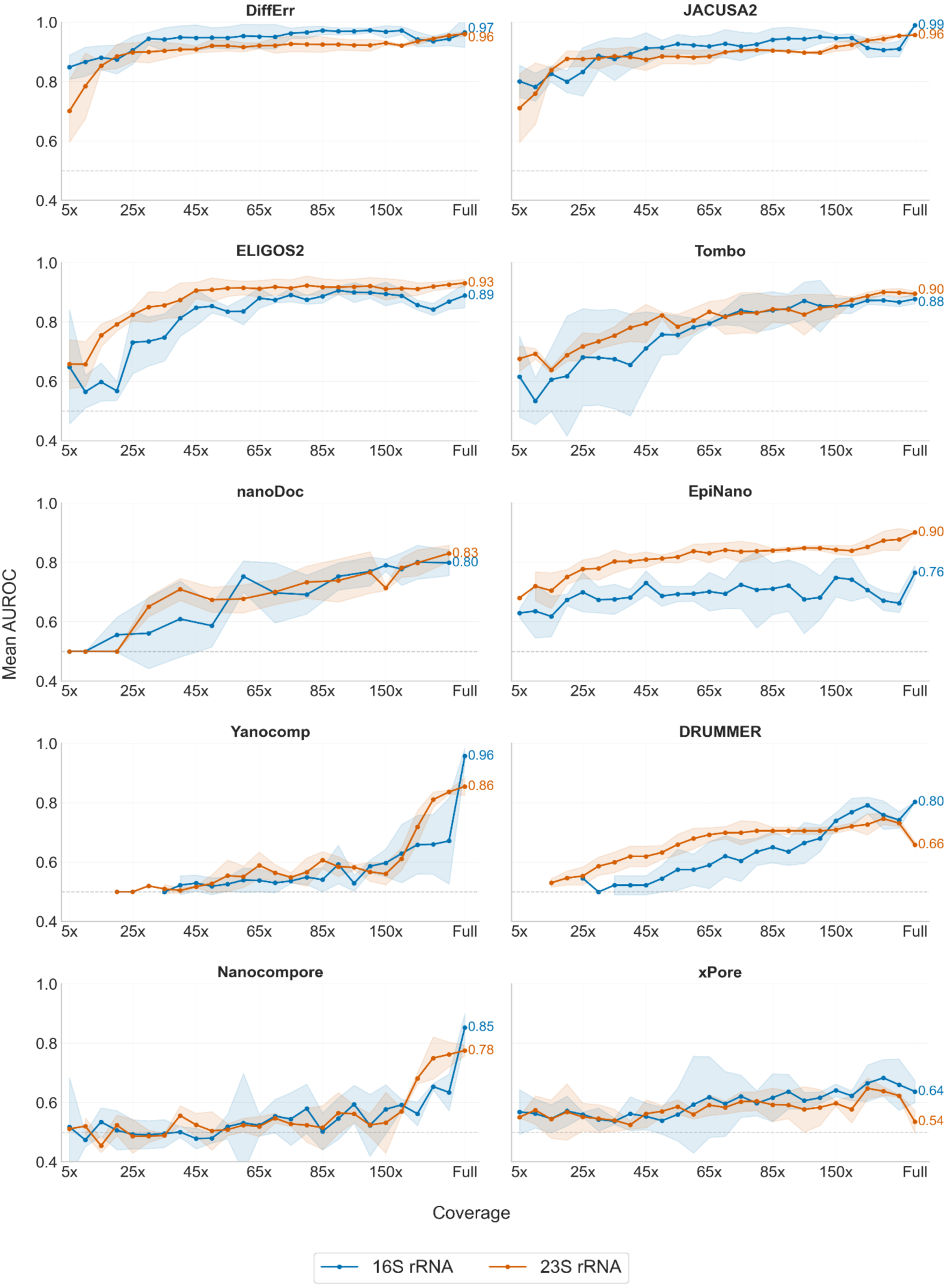
AUROC as a function of coverage across ten RNA modification detection tools. Each panel shows one tool, with separate curves for 16S rRNA (blue) and 23S rRNA (orange) across 25 coverage levels (5× to 1000×). Points represent mean AUROC across biological replicates; shaded bands indicate ±SD. The dashed line marks the random-classifier baseline (AUROC = 0.5). Each point indicates a 5x increase in coverage until 100×, after which the points are: 150×, 200×, 500×, 750×, and 1000× (with coverage only for alternating values in the case of nanoDoc).

As expected, AUROC increased monotonically with coverage for all tools in the subsampled data. (**Fig. 3**; **Supp. Fig. S3**). Only three tools (DiffErr, JACUSA2, and ELIGOS2) exceeded an AUROC of 0.9 within the subsampled coverage range (5×-1000×), with DiffErr requiring the least coverage on both rRNA molecules (25× for 16S; 30× for 23S). The remaining six tools did not reach 0.9 at any subsampled coverage level, with AUROC at 1000× ranging from 0.623 to 0.877 (**Supp. Table S4**).

At the full dataset coverage without subsampling (mean coverage: native 11,252x and IVT 3,375x on 16S; native 39,150x and IVT 5,640x on 23S), AUROC increased further for most tools, with the largest gains on 16S observed for Yanocomp (+0.286), Nanocompore (+0.219), and EpiNano (+0.102). This suggests that these tools were coverage limited even at 1000×. However, DRUMMER and xPore both exhibited decreased AUROC at 1000× for 23S (-0.073 and -0.087, respectively); and xPore AUROC also decreased for 16S (-0.023) at 1000×.

Given the severe class imbalance in our dataset (modification prevalence of 11 sites out of 1,542 for 16S rRNA and 25 out of 2,904 for 23S rRNA), we also used area under the precision-recall curve (AUPRC) (**Fig. 4; Supp. Fig. S4**). This curve quantifies how well rare modified sites are identified, rather than AUROC, which quantifies how well modified sites are discriminated from unmodified sites. On both 16S and 23S rRNA, DiffErr and JACUSA2 were again top performers, achieving AUPRC values of 0.4-0.5 at 1000× coverage. DRUMMER and ELIGOS2 reached intermediate values (0.2-0.44), though with more variation across different coverage levels. All other tools remained below 0.2 at every coverage level tested, and most stayed below 0.1 (**Fig. 4**).

**Figure 4.**
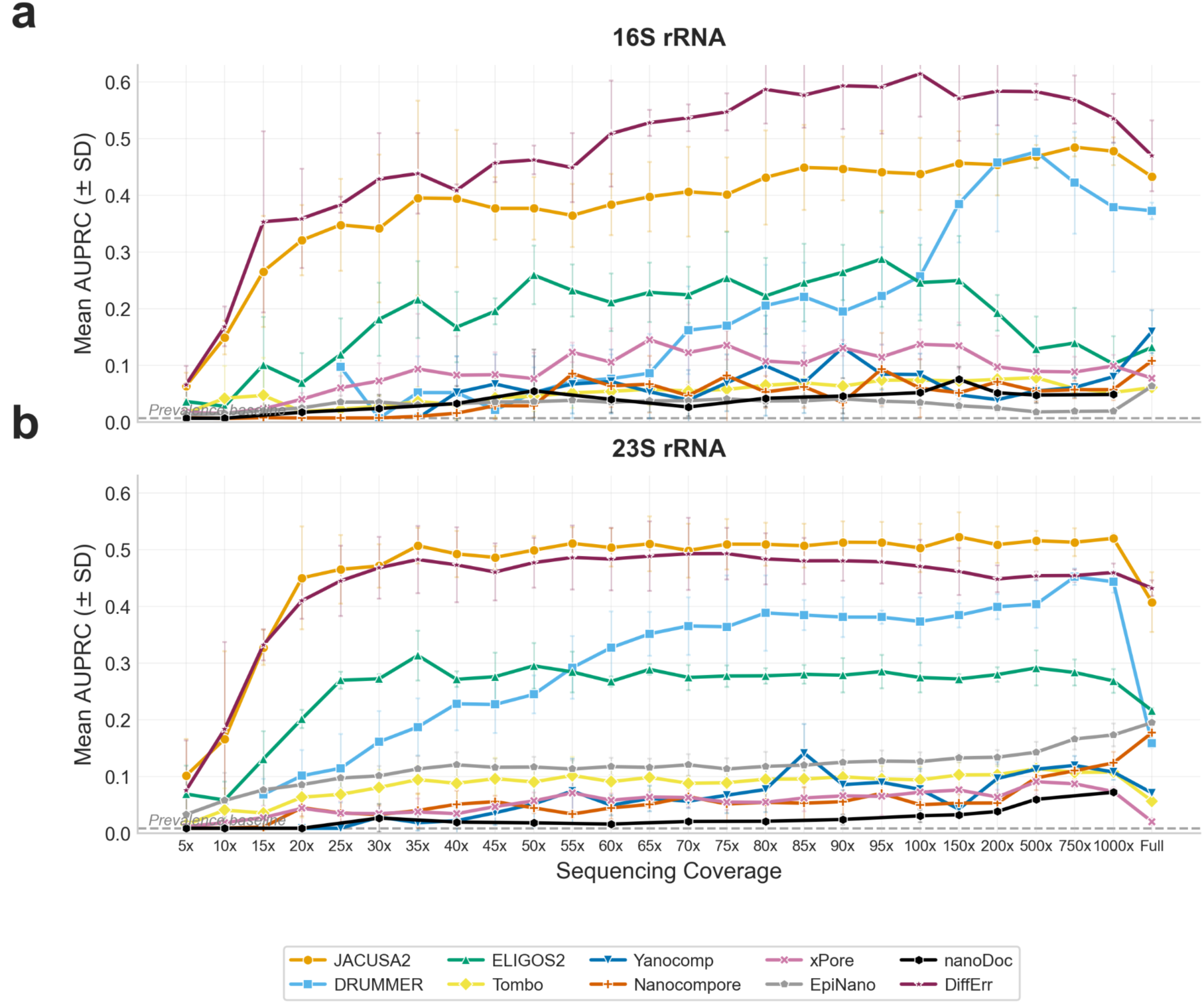
Precision-recall performance (AUPRC) across coverage levels. **(a)** 16S rRNA and **(b)** 23S rRNA across 25 coverage levels (5× to 1000×). Points represent mean AUPRC across biological replicates; error bars indicate ±SD. The dashed line marks the random-classifier baseline (equal to modification prevalence: ∼0.007 for 16S and ∼0.009 for 23S). The coverage levels plotted are the same as those in Fig. 3.

At the full dataset coverage, the differences between AUROC and AUPRC became more pronounced. While AUROC generally increased (as noted above), AUPRC declined for most of the top-performing tools (**Fig. 3 and 4)**. These patterns suggest that higher sequencing depths beyond 1000× can increase overall ranking accuracy (AUROC) while worsening precision-recall trade-offs (AUPRC), which may be driven by increased statistical power at high coverage, inflating false-positive calls at unmodified positions, and reducing precision without increasing true-positive recovery.

### Tool output completeness varies substantially and affects performance interpretation

The analyses presented above evaluate tools across all positions, assigning a score of zero to positions where a tool produces no output for that position. However, several tools did not emit scores for all positions, either due to the software coverage and/or read-count requirements; or because the models required a minimum number of reads to produce a result. To quantify this behavior, we measured two complementary metrics of output completeness: the overall call fraction (OCF), defined as the fraction of positions for which a tool produces any score, and the modified site call fraction (MCF), defined as the fraction of known modified sites that receive any score in the output, before applying any detection threshold (**Fig. 5**). Higher values of both metrics indicate more complete tool output. Tools exhibited distinct tiers of output completeness as measured by OCF and MCF (**Fig. 5a**). Four tools (Tombo, EpiNano, Nanocompore, and nanoDoc) maintained overall call fractions (OCF) at or near 100% across virtually all coverages on both rRNA species. JACUSA2, ELIGOS2, and DiffErr showed lower OCF at low coverages but converged to above 99.5% by 500×, placing them in the same high output completeness category at moderate-to-high depths. In contrast, DRUMMER and xPore maintained low OCF even at 1000× coverage: 6.6% and 29.5% on 16S rRNA and 6.2% and 32% on 23S rRNA, respectively. DRUMMER’s OCF remained below 7% at all coverage levels tested, indicating that its statistical model restricts output to a small, high-confidence subset of positions regardless of sequencing depth. xPore displayed a non-monotonic pattern, with its OCF clearly increasing from 29.4% at 25× to 42.2% at 100× before decreasing again to 29.5% at 1000×, consistent with increasingly stringent filtering at higher coverage. Yanocomp showed the strongest coverage dependence of any tool, with its OCF increasing from ∼1% at 50× to ∼78% at 1000×.

**Figure 5.**
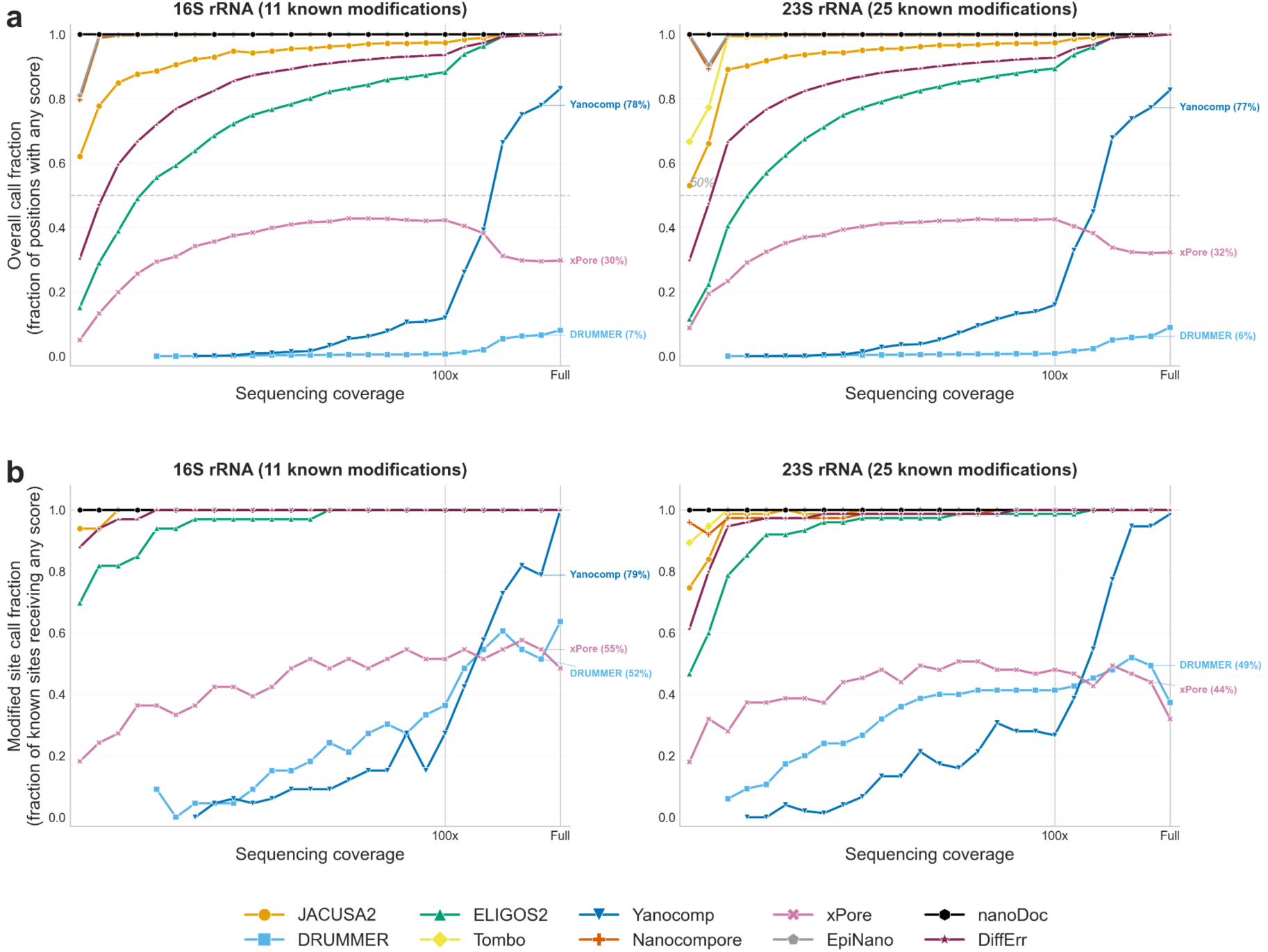
Output completeness metrics for ten tools evaluated across 25 different coverage levels (5-1000×) on *E. coli* 16S rRNA (11 known modifications, left) and 23S rRNA (25 known modifications, right). (**a**) Overall call fraction (OCF), defined as the fraction of all rRNA positions for which a tool produces any score. (**b**) Modified site call fraction (MCF), defined as the fraction of known modification sites that receive a score from the tool. This metric represents the theoretical maximum recall achievable by each tool; sites without scores cannot be detected regardless of threshold. The left panels show 16S rRNA and the right panels show 23S rRNA. Again, the coverage levels plotted are the same as those in Fig. 3).

These output-completeness patterns had direct consequences for the modified site call fraction (MCF) (**Fig. 5b**). Seven tools (Tombo, EpiNano, Nanocompore, JACUSA2, ELIGOS2, DiffErr, and nanoDoc) achieved 100% MCF at high sequencing depths (≥100× for most tools). At lower coverages, ELIGOS2, DiffErr, and JACUSA2 showed reduced MCF, while EpiNano and nanoDoc maintained 100% MCF across all coverages tested.

DRUMMER, by contrast, scored only approximately 6 of 11 known modification sites on 16S rRNA and 12 of 25 on 23S rRNA at 1000× coverage (MCF = 51.5% and 49.3%, respectively). Notably, DRUMMER’s MCF peaked at 500× (60.6% on 16S rRNA) and subsequently declined at 750× and 1000×, indicating that increased sequencing depth caused its model to become more selective, removing known modified sites, rather than more comprehensive. xPore showed a similar plateau, with MCF stabilising at approximately 54.5% on 16S and 44% on 23S across the 100-1000× range. Yanocomp reached near-perfect MCF at high depth (78.8% on 16S and 94.7% on 23S at 1000×), with only one modification site remaining unscored on each rRNA.

Using the full dataset without subsampling, OCF for the seven high-completeness tools converged to 100%. Among the selective tools, Yanocomp continued to improve, with its MCF reaching 100% on 16S and 98.7% on 23S, consistent with the large AUROC gains observed at full depth. By contrast, DRUMMER’s MCF declined on 23S (from 49.3% to 37.3%) and xPore’s declined on both rRNAs, indicating that the additional data caused these tools’ statistical models to become more restrictive. These drops in MCF provide a direct explanation for the AUPRC decreases observed for DRUMMER and xPore at full depth.

The practical consequence of selective reporting is that evaluation metrics can differ substantially depending on whether they are computed over all positions (with unreported positions assigned a score of 0) or only over positions where the tool produces a score (reported scope). For DRUMMER at 1000× on 16S rRNA, AUPRC was 0.379 (all positions) vs 0.719 (reported positions), a 1.9-fold increase that arises from excluding the 93.4% of positions on which DRUMMER reports no score. Similarly, xPore showed AUPRC of 0.099 (all) vs 0.178 (reported) on 16S, a 1.8-fold difference. This means that the performance is conditional on the subset of positions each tool evaluates. However, when comparing tools for their overall ability to identify modification sites across the transcript, evaluation across all positions is more appropriate because it incorporates both detection precision and the ability to provide scores locus-wide, and therefore does not favour tools that remain silent at many positions. All subsequent analyses evaluate across all positions, and we report both approaches in **Supplementary Table S7** to enable tool evaluation under either framework.

### Signal-based tools show systematic 5ʹ directional bias in modification localisation

To assess the positional precision of each tool, we evaluated detection performance at 21 directional offsets (δ = -10 to +10 nucleotides) from known modification sites at 1000× coverage. In this analysis, to quantify detection metrics, the ground truth known modification is shifted by δ nucleotides, such that positive δ values represent downstream (3ʹ) displacement and negative δ values represent upstream (5ʹ) displacement relative to the true modification site. DiffErr, JACUSA2 and DRUMMER showed sharp AUPRC peaks at the exact modification position (δ = 0), with values 0.536, 0.478 and 0.379 on 16S rRNA; 0.460, 0.520 and 0.443 on 23S rRNA, respectively. For these three tools, AUPRC at neighbouring offsets dropped to near-baseline levels, indicating precise single-nucleotide localisation.

In contrast, the remaining tools exhibited AUPRC profiles that peaked at negative offsets, indicating systematic displacement of the detected signal toward the 5ʹ direction, though the magnitude of displacement varied (**Fig. 6b**). For 16S rRNA, Tombo and nanoDoc showed largest offsets, both peaking at δ = -4 (AUPRC = 0.185 and 0.273, respectively) compared to substantially lower values at δ = 0 (0.052 and 0.049). For 23S rRNA, nanoDoc again peaked at δ = -4 (AUPRC = 0.237 vs 0.072 at δ = 0), while Tombo peaked at δ = -2 (-0.151 vs 0.107 at δ = 0). Nanocompore peaked at δ = -2 on 16S (AUPRC = 0.219 vs 0.057 at δ = 0) and δ = -1 on 23S (0.218 vs 0.124). EpiNano showed the largest improvement at its offset relative to the centred δ = 0 position: peak AUPRC at δ = -1 was 0.408 on 16S and 0.505 on 23S, compared to 0.020 and 0.173 at δ = 0, indicating that the majority of its detection signal falls one nucleotide upstream of the annotated site. ELIGOS2 and Yanocomp also peaked at δ = -1 on 16S (AUPRC = 0.164 and 0.138, respectively), though ELIGOS2 peaked at δ = 0 on 23S (0.268) while Yanocomp peaked at δ = -1 on both rRNAs. xPore showed an inconsistent pattern, peaking at δ = -1 on 16S (0.107) but δ = +1 on 23S (0.109) (**Fig. 6b**).

**Figure 6.**
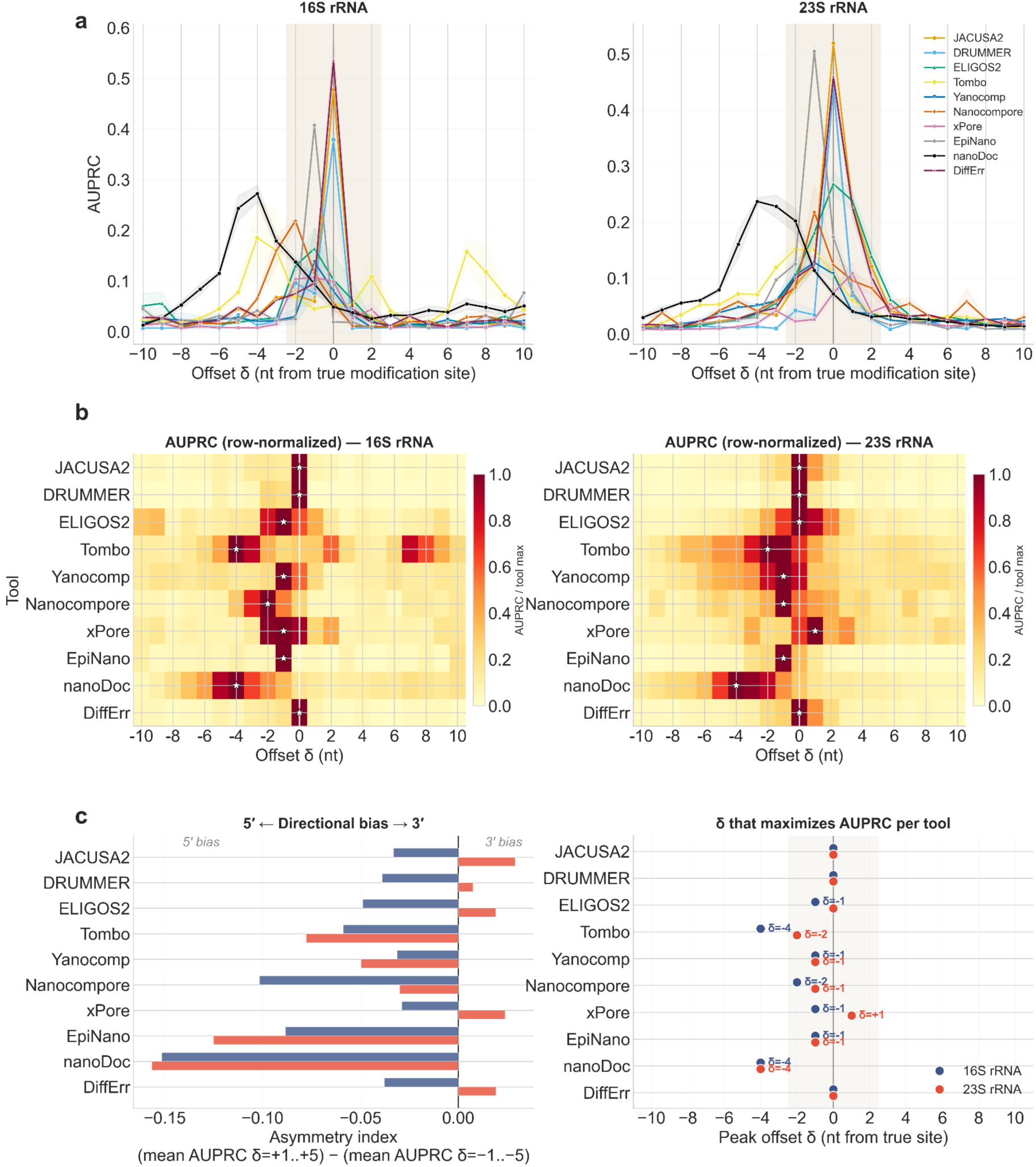
Directional offset analysis reveals systematic 5ʹ localisation bias in signal-based modification detection tools. Performance of ten tools evaluated at 21 directional offsets (δ = -10 to +10 nt) from known modification sites on *E. coli* 16S and 23S rRNA at 1000× coverage. Negative δ values indicate upstream (5ʹ) displacement; positive δ values indicate downstream (3ʹ) displacement. **(a)** AUPRC across directional offset δ for each tool. Shaded bands indicate ±1 SD across replicates. The beige region highlights the ±2 nt range corresponding to the approximate 5-mer reader head context. **(b)** Row-normalised AUPRC heatmaps (each tool’s value scaled to its own maximum). White stars mark the peak offset per tool; the white vertical line indicates δ = 0 (true modification position). **(c)** Left: asymmetry index, defined as the difference between mean AUPRC over δ = +1 to +5 and mean AUPRC over δ = -1 to -5. Negative values indicate a 5ʹ bias (modification signal detected upstream of the true site). Right: peak offset δ that maximises AUPRC for each tool, shown separately for 16S (blue) and 23S (red) rRNA. Tools that peak at δ = 0 achieve exact positional localisation; tools peaking at negative δ values systematically mislocalise modification signals towards the 5ʹ direction.

To quantify directional asymmetry, which captures bias in the spread of the signal peak versus the maximum value of the signal peak, we computed an asymmetry index. We defined this as the difference between mean AUPRC over δ = +1 to +5 and mean AUPRC over δ = -1 to -5, such that negative values indicate 5ʹ bias (**Fig. 6c** left panel). All signal-based tools showed consistently negative asymmetry indices across both rRNA molecules. nanoDoc exhibited the strongest 5ʹ bias (-0.152 on 16S; -0.157 on 23S) followed by EpiNano (-0.088 on 16S; -0.125 on 23S), Nanocompore (-0.102 on 16S; -0.030 on 23S), and Tombo (-0.059 on 16; -0.078 on 23S). DiffErr, JACUSA2, and DRUMMER showed low asymmetry on both rRNAs (absolute values ≤0.039), consistent with their peak values at δ = 0. Most tools showed reproducible directional bias for both rRNA molecules (**Fig. 6c** right panel): DiffErr, JACUSA2, DRUMMER, nanoDoc, EpiNano, and Yanocomp peaked at the same offset on both 16S and 23S. However, Tombo shifted from δ = -4 (16S) to δ = -2 (23S), Nanocompore from δ = -2 to δ = -1; most surprisingly, xPore peaked at δ = -1 on 16S but δ = +1 on 23S.

Consistent with these offset patterns, recomputing AUPRC with symmetric tolerance windows of ±1 to ±10 nucleotides, as used in previous benchmarks (Jenjaroenpun et al., 2021; Spangenberg et al., 2025), shifted tools rankings in favour of displaced tools and penalised those with precise localisation (**Supp. Fig. S9**). However, because several modification sites on *E. coli* rRNA are separated by fewer than 10 nucleotides, larger tolerance windows can progressively merge distinct ground-truth sites into overlapping zones, inflating apparent modification prevalence up to 13-fold at ±10 and making resulting AUPRC values difficult to interpret.

### Tools with strong ranking performance can still produce many false positives at usable thresholds

To complement the AUROC and AUPRC analyses and assess the practical operating characteristics of each tool, we examined performance at each tool’s maximal F1 threshold using 1000× coverage data (no offset; see analysis below), where most tools had reached high or near peak performance. In precision-recall space (**Fig. 7**), JACUSA2 showed a favourable balance on 16S rRNA, combining high precision (0.89) with moderate recall (0.45) while producing fewer than one false positive on average out of 1,531 unmodified positions. DiffErr achieved the highest F1 score on 16S (0.68), combining high precision (0.77) with the joint-highest recall (0.61) and approximately 2 false positives. DRUMMER achieved a similar F1 score to JACUSA2 (0.6) with slightly higher recall (0.52) but at lower precision (0.71) and approximately 2.3 false positives. By contrast, Tombo reached the highest recall (0.61) but with a substantially larger false-positive burden of 92 false positives (**Fig. 8**). On 23S rRNA, which has a broader diversity of modification types, DRUMMER had the highest precision (0.81) with only 2.7 false positives out of 2,879 unmodified positions at recall 0.45, while JACUSA2 recovered more modification sites (recall 0.59) at reduced precision (0.48) and approximately 16 false positives. These results were broadly consistent with ranking-based metrics: DiffErr and JACUSA2 were the strongest performers on 16S, while DRUMMER achieved the highest precision and F1 on 23S rRNA. Importantly, these metrics should be interpreted alongside the OCF (**Fig. 5**), since tools with low OCF, such as DRUMMER and xPore, may appear more precise partly because they evaluate fewer positions.

**Figure 7.**
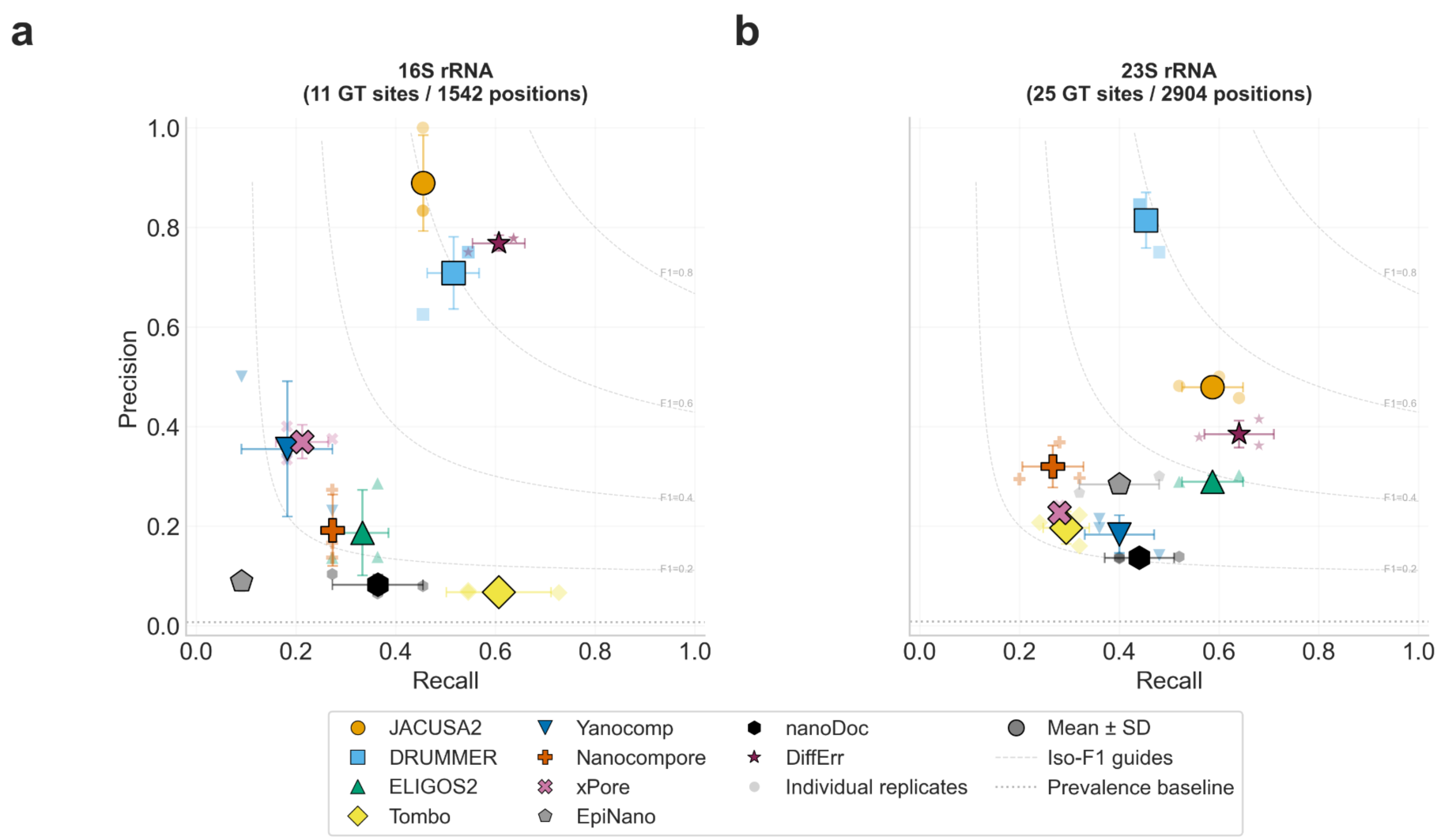
Precision-recall operating points for all tools at 1000× coverage. Each tool is plotted at its maximal F1 threshold for 16S rRNA (a) and 23S rRNA (b). Large symbols show the mean across three replicates; GT = Ground Truth (i.e. known modified sites); small symbols show individual replicates. Error bars indicate standard deviation. Dashed grey curves are iso-F1 contours (lines of equal F1 score). The dotted horizontal line marks the prevalence baseline (proportion of true modifications to total positions in each rRNA), corresponding to the expected precision of a random classifier. The ideal operating region is the upper-right corner (high precision, high recall).

**Figure 8.**
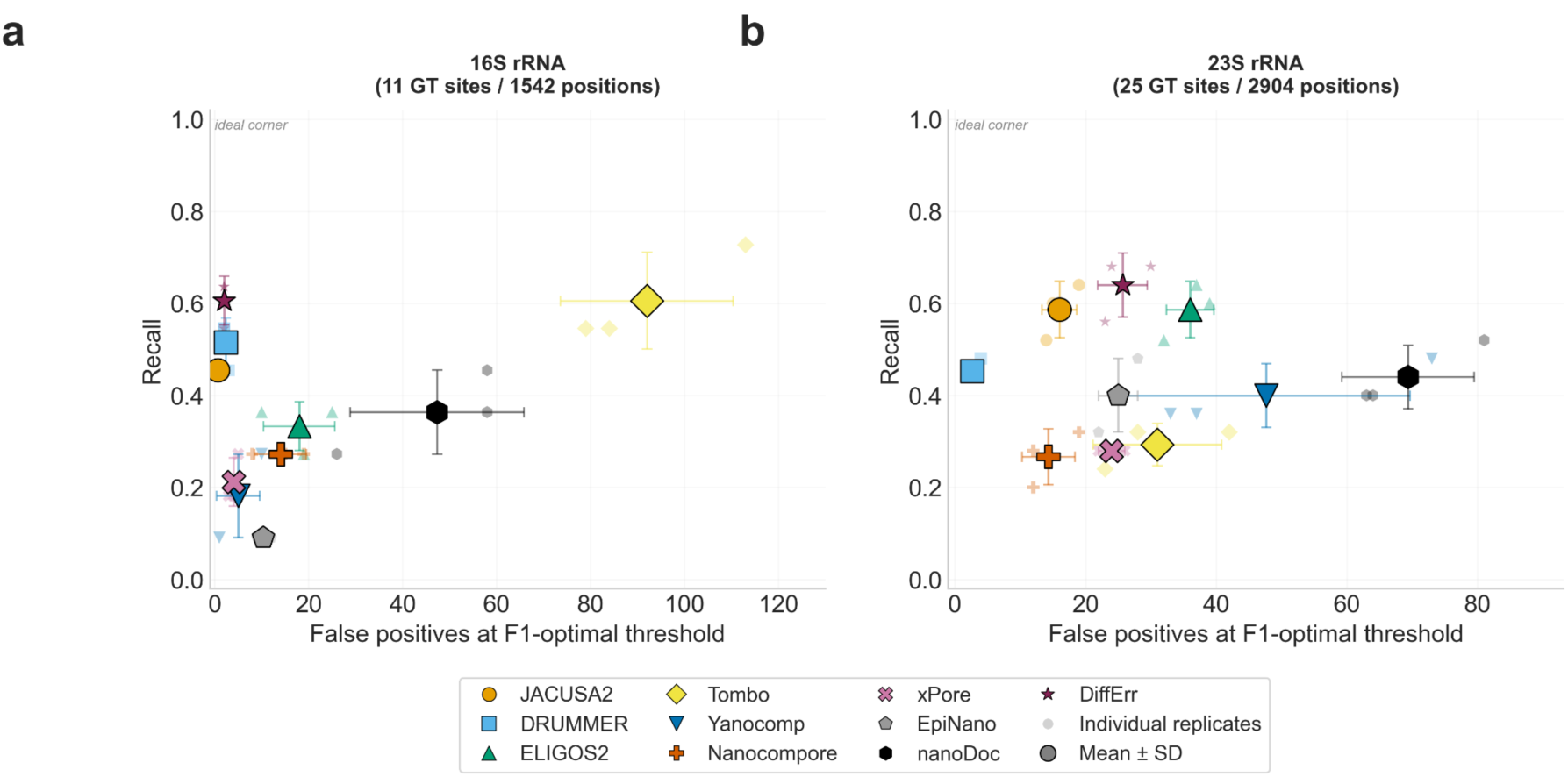
Recall versus false-positive count at maximal F1 thresholds for all tools at 1000× coverage. Each tool is shown for 16S rRNA (**a**) and 23S rRNA (**b**). The x-axis shows the absolute number of false-positive calls; the y-axis shows recall (fraction of ground-truth sites recovered). Large symbols show mean across three replicates; small symbols show individual replicates. Error bars indicate standard deviation. The ideal operating region is the upper-left corner (high recall, few false positives).

### Detection of individual modification sites is strongly dependent on both modification type and tool

To understand which specific modification sites drive the recall differences observed in aggregate metrics, we examined detection at each of the 11 (16S rRNA) and 25 (23S rRNA) known ground-truth (modification) sites across all tools and replicates at 1000× coverage with no offset correction (see below) (**Fig. 9**; per replicate data is in **Supp. Fig. S5-S6**). This analysis revealed that no single site on 16S rRNA was consistently detected (3/3 replicates) by all tools, and the most broadly detected sites (516 Ψ, 1207 m2G) were each recovered by only 5 of 10 tools. On 23S rRNA, only three sites (2069 m7G, 2503 m2A, and 2604 Ψ) were detected by seven or more tools. Tool-specific detection patterns were non-overlapping; on 16S, DiffErr detected the most sites consistently (6/11 for 3/3 replicates), followed by JACUSA2, DRUMMER and Tombo (each 5/11 sites). However, the five sites detected differed partly between tools. Tombo additionally detected four sites in only one or two replicates, indicating lower reproducibility at the per-site level. JACUSA2 showed the most reproducible pattern: every site it detected was detected in all three replicates, with no partial calls.

**Figure 9.**
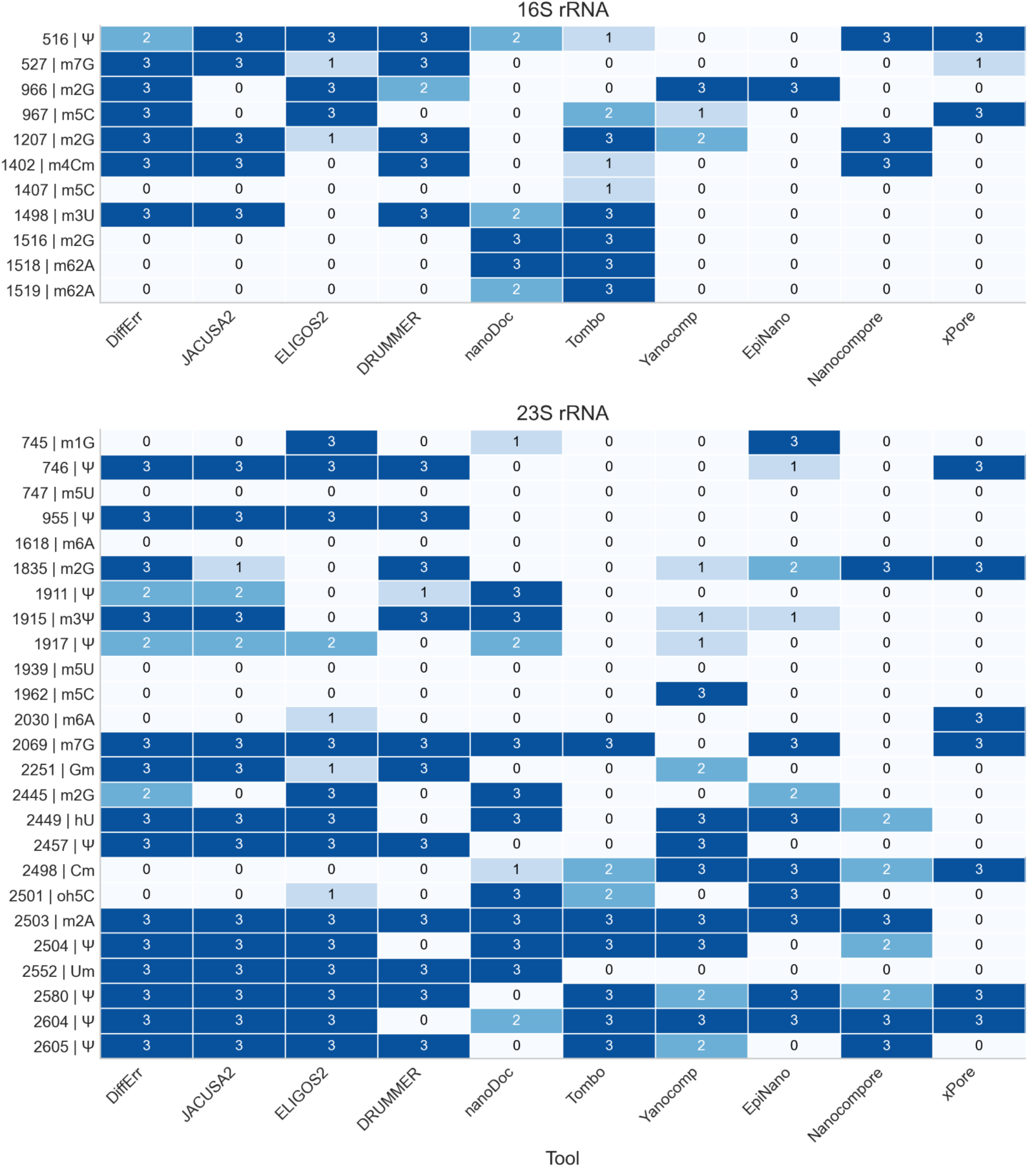
Ground-truth (modification) site recovery across tools at 1000× coverage. Each cell shows the fraction of replicates (out of 3) in which a known modification site was detected at that tool’s maximum F1 score, for 16S rRNA (**top**) and 23S rRNA (**bottom**). Darker shading indicates detection in more replicates. Sites are ordered by genomics position (rows), tools are ordered by overall detection rate (columns). Site labels indicate position and modification type. Per replicate detection patterns shown in **Supp. Figures S4** and **S5**.

Several modification types proved particularly challenging. The two m5C sites on 16S showed divergent detectability: position 967 was recovered by three tools consistently (DiffErr, ELIGOS2, xPore) with two additional tools detecting it in a subset of replicates, while position 1407 was missed by all tools except Tombo (1/3 replicates). For 23S, both m5U sites (747, 1939) were undetected by all tools. The m5C site at position 1962 was detected only by Yanocomp (3/3 replicates). Similarly, both m6A sites on 23S (1618, 2030) were largely invisible, with only xPore recovering position 2030 in all three replicates.

DRUMMER’s high precision (**Fig. 7 and Fig. 8**) may partly be explained by its conservative reporting model (i.e., its Bonferroni correction, which restricts output to only the highest-confidence sites). This restricts the output to a small, high-confidence subset, limiting false positives but also reducing recall. Five of its six undetected ground-truth sites on 16S were positions for which DRUMMER produced no output, rather than positions scored below threshold.

### Offset correction reveals underestimated detection capacity in mislocalised tools

The per-site analysis above and the precision-recall operating points in the preceding sections both evaluate tools at the exact annotated modification position (δ =0). However, the directional offset analysis (**Fig. 6**) established that several tools systematically detect modification signals 1-4 nucleotides upstream of the true site. This raises the question of whether the sites apparently missed by these tools are simply mislocalised. To address this, we re-evaluated each tool’s precision, recall, and site-level recovery at its empirically determined peak offset rather than at δ = 0 (**Supp. Tables S5-S6**).

Tools with precise localisation (DiffErr, JACUSA2, DRUMMER) showed no change, as expected. However, offset correction substantially improved the operating characteristics of several displaced tools, both by recovering additional modification sites and by reducing false positives. Improvement for EpiNano was extremely high: with a single nucleotide shift (δ = -1), its F1 Score increased from 0.090 to 0.515 on 16S and 0.331 to 0.560 on 23S. For 16S, EpiNano recovered 6 of 11 ground-truth sites at its peak offset compared to only 1 at δ = 0, while simultaneously reducing false positives from 10 to 6. On 23S, it gained approximately 2 additional sites while false positives dropped from 25 to 5. nanoDoc similarly improved on 16S, recovering 7.3 sites (vs 4.0 at δ = 0) while also reducing false positives from 45 to 21. For Tombo, site recovery remained similar (6.7 to 6.0) but false positives dropped from 92 to 19, turning from practically unusable to interpretable.

These results indicate that a substantial fraction of the performance between error-rate based tools (DiffErr, JACUSA2, DRUMMER) and signal-based tools (EpiNano, Tombo, nanoDoc) is attributable to positional mislocalisation rather than an inability to detect modifications. The offset estimates reported here (**Supp. Tables S5-S6**) can serve as a practical guide for this correction, but may differ for other RNA species or modification types.

### Tool combinations improve site recovery

Given the non-overlapping detection patterns across tools, we then examined whether combining outputs from tools could improve site recovery while controlling false-positive burden. For each combination of one, two, or three tools, we calculated the union of positive calls across both rRNAs and quantified the resulting true positives (ground-truth sites detected in ≥2/3 replicates) and false positives (**Fig. 10**).

**Figure 10.**
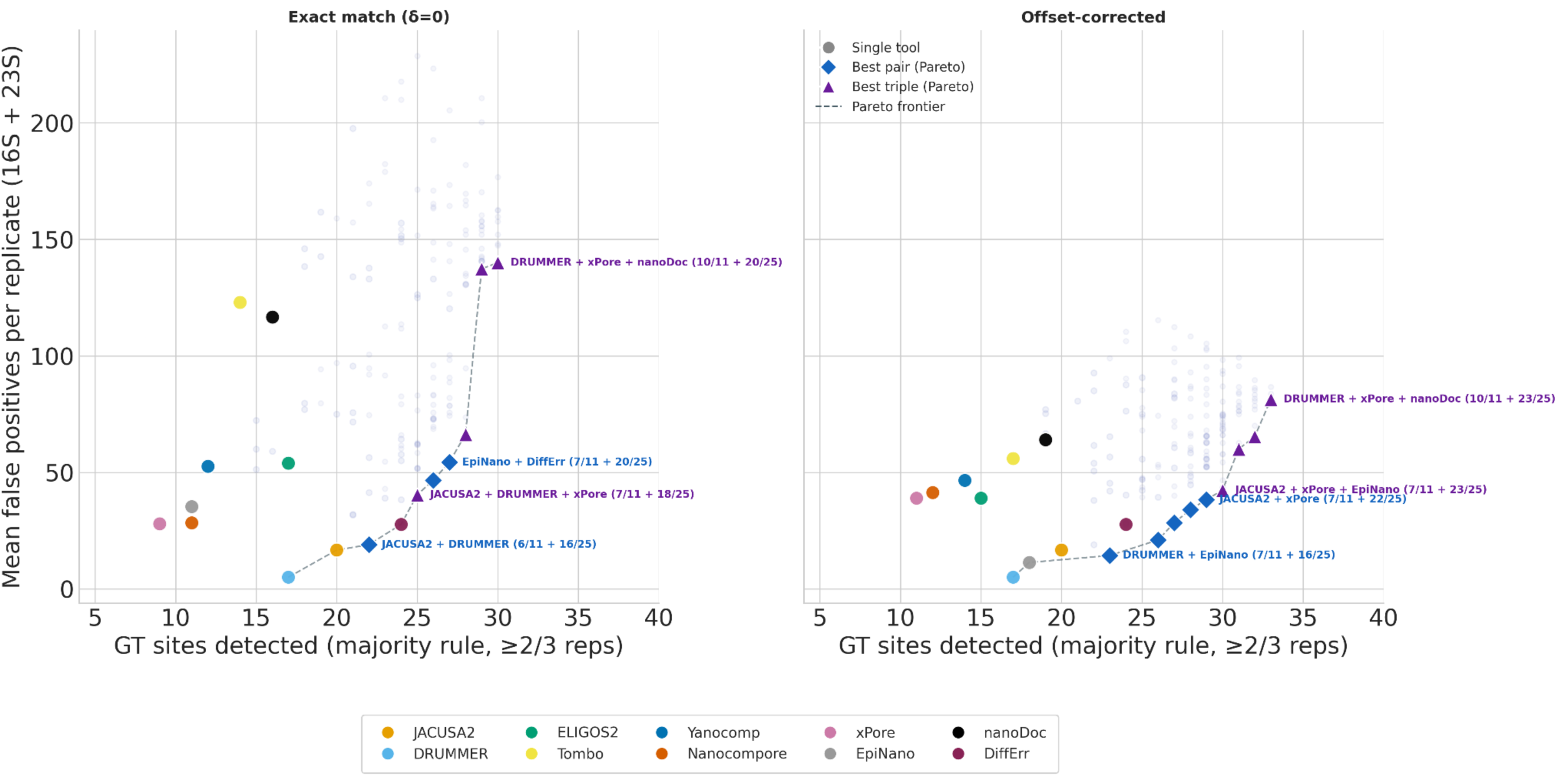
Tool combinations for ground-truth site recovery at 1000× coverage. Each point represents a single tool (coloured circles), the best Pareto-optimal pair (blue diamonds), or the best Pareto-optimal triple (purple triangles). The x-axis shows the number of ground-truth modification sites detected in ≥2 of 3 replicates (out of 36 total: 11 on 16S + 25 on 23S rRNA). The y-axis shows the mean number of false positives per replicate (summed across both rRNAs). Faint dots show all possible pair and triple combinations. The dashed line traces the Pareto frontier, where no combination detects more sites without increasing false positives. Left panel: exact-position evaluation (δ=0). Right panel: offset-corrected evaluation, where each tool’s score is assessed at its empirically determined peak offset. The Pareto frontier shifts rightward after offset correction, indicating that more ground-truth sites are recoverable. Key Pareto-optimal combinations are labelled with their 16S and 23S site recovery.

At exact-position evaluation (δ=0), the highest precision single tool was DRUMMER (17/36 sites, FP=5), and the best pair was JACUSA2 + DRUMMER (22/36 sites, FP=19). Reaching 30/36 sites required a triple tool combination (DRUMMER + xPore + nanoDoc) at a cost of ∼140 false positives. After applying tool-specific offset corrections, the Pareto frontier shifted substantially towards right: DRUMMER + EpiNano recovered 23/36 sites with only 14 false positives, and JACUSA2 + xPore + EpiNano recovered 30/36 sites (7/11 on 16S + 23/25 on 23S) with 42 false positives, a three fold reduction in FP compared to equivalent δ=0 triple tool combination. The best offset-corrected triple (DRUMMER + xPore + nanoDoc) recovered 33/36 sites (92%) with ∼81 false positives (**Supp. Table S8**).

Five modification sites remained undetected by any tool or combination even after offset correction: m5C at position 1407 on 16S, and on 23S both m5U sites (747, 1939), m6A (1618), and pseudouridine (1917). These modification types seem particularly difficult for current DRS-based comparative approaches to reliably detect on *E. coli* rRNA.

## Discussion

RNA modification detection from nanopore direct RNA sequencing (DRS) remains a moving target because tool performance depends not only on the underlying algorithm but also on sequencing chemistry, training data, type of modification, sequence context, coverage, and the availability of matched controls. In this context, our benchmark offers a unique and complementary perspective by evaluating ten comparative computational tools on a densely modified, biologically well-annotated bacterial model: *Escherichia coli* K-12 MG1655 16S and 23S rRNA, which together contain 36 known modified sites spanning 17 modification types. While most previous benchmarks have focused on m6A detection in eukaryotic mRNA, typically from human or mouse datasets (Zhong et al., 2023b; Maestri et al., 2024; Zou et al., 2025; Ohnezeit et al., 2026), our evaluation covers multi-modification type detection in bacterial rRNA using native RNA and matched IVT controls across 25 coverage levels and three biological replicates. This design reveals aspects of tool behaviour that conventional summary metrics alone do not capture, particularly output completeness (overall call fraction, OCF), positional precision, and per-site recoverability across diverse modification chemistries, as well as sensitivity and precision increases upon implementation of optimal offsets.

DiffErr and JACUSA2 emerged as the strongest overall performers in our benchmark. Differr attained the highest AUROC on 16S rRNA (0.944) and tied with JACUSA2 on 23S (both 0.955), while also achieving the highest F1 score on 16S (0.68) with high precision and the joint-highest recall. JACUSA2 showed the most consistent precision-recall balance across both rRNAs, the most reproducible per-site detections, and together with DiffErr and DRUMMER, precise single-nucleotide localisation at the true modification position. Notably, JACUSA2 (Piechotta et al., 2022) has not been included in any of the major published tool benchmarks to date. In those studies, m6Anet consistently emerged as a top performer for m6A detection (Zhong et al., 2023b; Maestri et al., 2024; Luo et al., 2026). Comparative tools like ELIGOS and Nanocompore have been previously shown as performing best in yeast (Maestri et al., 2024). However, these benchmarks focused on m6A in eukaryotic mRNA, where deep-learning tools (e.g. m6Anet) that are trained on mammalian DRACH motifs have an inherent advantage. In bacterial rRNA, where modification types are diverse and lie outside canonical eukaryotic motifs, error-comparison frameworks that are not reliant on pretrained modification-specific models performed better than signal-based and deep-learning tools. These results indicate that tool rankings are substrate-specific, and that top performers from eukaryotic m6A benchmarks should not be assumed to generalise to bacterial or multi-modification settings.

The relationship between sequencing coverage and tool performance proved more nuanced than expected. AUROC increased with coverage for all tools but the rate of improvement and point of stabilisation varied: DiffErr and JACUSA2 reached near-peak AUROC by 100× on 16S, where as tools with high no-call rates (Yanocomp, Nanocompore) continued improving up to 1000× and beyond. AUPRC showed a different and arguably more informative pattern: for most tools, AUPRC peaked at moderate coverages (50-100×) and declined at higher depths, particularly in the full unsubsampled data. This divergence between AUROC and AUPRC at high coverage likely reflects an increasing false-positive burden, perhaps due to increased statistical power at high coverage, that decreases precision without proportionally improving true-positive recovery. Our results suggest that ≥ 100× coverage is sufficient for the top performing tools (DiffErr and JACUSA2), while tools with low OCF require ≥500× to approach their performance ceiling.

Output completeness represents a dimension of tool performance that is rarely reported but substantially affects interpretation. We found that the tools differ dramatically in how many positions they actually score, and that this directly inflates apparent performance when evaluation is restricted to reported positions. For example, DRUMMER is very conservative and reports scores for only ∼6% of positions even at 1000× coverage, which led to high precision within that subset but missed approximately half of all ground-truth sites. Previous benchmarks have noted related problems without measuring them directly. For example, (Maestri et al., 2024) reported difficulties running several tools and the generation of empty output files, and (Tan, Guo, Wang, et al., 2024b) found that DRUMMER and EpiNano_Error produced very few to no outputs on viral RNA. Our results go further by quantifying how these incomplete outputs inflate tools’ apparent accuracies and establishing that OCF and MCF should be reported alongside discrimination metrics as standard benchmarking practice.

There are at least two practically important findings of our study. The first is the systematic 5’ positional offset exhibited by many signal-based tools. Positional offsets arise due to the nanopore read head containing approximately five nucleotides at a time as RNA translocates through the pore, and we found that tools operating on raw current signals (Tombo, Nanocompore, nanoDoc, EpiNano) consistently showed their strongest signal 1-4 nucleotides upstream of the known modified site. Error-rate-based tools (DiffErr, JACUSA2, DRUMMER) localised accurately at the correct position. Previous studies have accommodated potential positional imprecision by using tolerance windows, such as ±1 nucleotide (Spangenberg et al., 2025), but none have characterised the direction, magnitude, or tool-specificity of the offset. Independently, (Riquelme-Barrios et al., 2025) observed a 5-nucleotide upstream offset for the m3U modification at position 1498 of *E. coli* 16S rRNA using ELIGOS, supporting the systematic 5’ directional bias we characterise here across multiple tools.

Critically, we also showed that applying tool-specific offset corrections transformed the performance of several tools. EpiNano, ranked lowest at exact-position evaluation, became one of the strongest performers with a single-nucleotide correction (F1: 0.09 to 0.52 on 16S), recovering five additional ground-truth sites while also reducing false positives. Site-level combination analysis confirmed these improvements: with offset correction, DRUMMER + EpiNano recovered 23 of 36 sites with only 14 false positives, whereas the equivalent pair at δ=0 (JACUSA2 + DRUMMER) recovered 22 sites with 19 false positives. These results indicate that a substantial fraction of the performance gap between error-rate-based and signal-based tools is attributable to mislocalisation rather than inability to detect modifications, and that best practices for signal-based tools should include the application of offset corrections. While not always available, in many cases these can be inferred when ground truth data is available, which may be a frequent occurrence.

Second, we quantified the increases in sensitivity and precision that arise via tool combination. We showed that with a known ground truth, these combinations can be systematically tuned along the Pareto front to maximise desired goals (e.g. sensitivity).

Importantly, aggregate metrics can also mask variation in per-site detectability, and our site-level analysis revealed that tools often fail on specific modification classes. Most m5C, m5U, and m6A sites on both rRNA molecules were undetected by majority of tools, while only three 23S sites: 2069 m7G, 2503 m2A, and 2604 Ψ were recovered by seven or more tools. These site-specific detection patterns are consistent with another study (Riquelme-Barrios et al., 2025) which failed to detect m5C at positions 967/1407 (16S) and 1962 (23S), and m6A at position 1618 (23S) using ELIGOS. Low inter-tool concordance of sites identified has also been observed in eukaryotic m6A benchmarks, such as (Luo et al., 2026), who found overlap between any two tools was typically below 50%. The poor recovery of m6A sites is particularly notable because m6A is the best-performing modification in eukaryotic benchmarks. One possible explanation is that *E. coli* rRNA m6A sites lie outside the canonical DRACH motifs that dominate training data for most eukaryotic-centric m6A tools, consistent with the distinct GACGCMAG-enriched sequence contexts reported for *E. coli* m6A (Tan, Guo, Shao, et al., 2024). More generally, our findings agree with previous work showing that detectability depends strongly on modification type, and sequence context (White et al., 2024; Spangenberg et al., 2025; Luo et al., 2026).

There are some limitations in this study. First, the benchmark was performed on bacterial rRNA, which may have systematically different signal patterns compared to mRNA, for example due to folding or changes in translocation speed (dwell time), which is used by some tools as information site modification probabilities (Leger et al., 2021). Second, modifications on *E. coli* rRNA are extremely consistent, being present on nearly every RNA molecule, with over 90% of the copies carrying each modification under standard growth conditions (Popova & Williamson, 2014). Our benchmark tests detection of modifications that are almost always present, in contrast to mRNA, for which modifications may occur on only a fraction of molecules. Thus, the results here likely represent a best-case scenario for detection sensitivity. Third, all data were generated with RNA002 kit chemistry (R9.4.1 pores). ONT has since released the RNA004 kit with RNA-specific pore chemistry. However, it is not clear that RNA004 tools consistently outperform RNA002 tools for non-m6A modifications (Luo et al., 2026), and that even with retraining, non-m6A tools can continue to show low AUPRC and limited precision. Minimally though, optimal tool parameters, OCF, and positional offsets may differ across sequencing chemistries and would need re-evaluation. Finally basecaller-integrated methods (e.g., Dorado modified basecalling, m6Anet) were outside the scope of this benchmark, and their single-molecule sensitivity could provide a more complete picture of the current tool landscape in bacterial context.

Taken together, our results establish that discrimination metrics alone are insufficient to evaluate RNA modification detection tools. We show that output completeness, positional precision, and per-modification-type sensitivity are critical metrics alongside AUROC and AUPRC as standard benchmarking outputs. We demonstrate that signal-based tools carry systematic 5’ positional offset that, once characterised with ground truth, can be corrected to substantially improve per-site recovery. We also show that sequencing coverage of at least 100× is sufficient for top-performing tools to reach near-peak discrimination precision, though higher depths are needed for some tools. Our combination analysis demonstrates that pairing a precisely localised error-rate tool with an offset-corrected signal-based tool consistently outperforms same-category combinations. Despite this, certain modification types, particularly m5C, m5U, and m6A, are systematically missed by almost all tools. This may reflect the eukaryotic m6A-centric training data that underlies most current methods. Future work should assess where these findings generalise to data from RNA004 DRS kit chemistry, developing organism-specific training datasets and retraining frameworks to improve detection of these under-represented modifications.

## Methods

### RNA Isolation

We inoculated three colonies of *E. coli* K12 MG1655 into 5 mL of M9 minimal media with 0.2% glucose and incubated overnight in a shaking incubator at 37°C. We diluted the overnight cultures 1:50 into fresh 25 mL M9 minimal media with 0.2% glucose and grew them at 37°C to mid-exponential phase (OD 0.5-0.6). To stabilise RNA, we added 0.5 volumes of ice cold 5% phenol in ethanol and incubated on ice for 15 minutes. We pelleted the bacteria in a centrifuge at 7000 x g for 7 minutes at 4°C, resuspended in 350 µL lysozyme solution (2 mg/mL), and incubated for 5 minutes. An equal volume (350 µL) of RNA lysis buffer (Zymo Quick RNA miniprep kit, cat# R1054) was added and mixed thoroughly. Following this, we isolated the total RNA using the manufacturer’s protocol and eluted in 100 µL of RNAse-free water.

### RNA Poly(A) Tailing

We polyadenylated total RNA to enable direct RNA sequencing library preparation using the *E. coli* poly(A) polymerase kit (New England Biolabs, cat# M0276). Briefly, we incubated 2 µg of total RNA with 2 µL of 10x reaction buffer, 2 µL of 10 mM ATP, and 1.5 µL of poly(A) polymerase in a total reaction volume of 20 µL (volume made up with nuclease-free water) at 37°C for 30 minutes. Polyadenylated RNA was then purified using AMPure RNA XP beads (Beckman Coulter, cat# A66514) following the manufacturers protocol and eluted in nuclease-free water.

### *In vitro* Preparation of Unmodified RNA

To produce unmodified RNA as a negative control for modification detection, we performed transcriptome-wide *in vitro* transcription (IVT) as described previously (protocols.io: https://www.protocols.io/view/synthesis-of-in-vitro-transcribed-rna-from-whole-b-81wgb7r7yvpk/v1). Briefly, we generated full-length cDNA from polyadenylated *E. coli* total RNA using a modified template-switching approach with Moloney Murine Leukemia Virus Reverse Transcriptase (MMLV RT). We used a custom strand-switching primer containing the T7 promoter sequence at the 5ʹ end (**Supp. Table. S3**) to enable T7-mediated transcription from the resulting cDNA, yielding full-length double-stranded cDNA templates with a 5ʹ T7 promoter. IVT RNA produced in this manner contains only canonical (unmodified) bases, making it suitable as a matched negative control for comparative modification detection.

### Barcoding and Demultiplexing

At the time of this study, commercially available Oxford Nanopore direct RNA sequencing kits did not support multiplexing. We therefore used custom barcodes for multiplexing and a demultiplexing tool (Deeplexicon) as described in (Smith et al., 2020).

### Direct RNA library preparation and sequencing

We prepared direct RNA sequencing libraries using the Oxford Nanopore Technologies SQK-RNA002 kit according to the manufacturer’s protocol. Sequencing was performed on a MinION device using R9.4.1 flow cells.

## Data Analysis

### Basecalling

We basecalled FAST5 files obtained from sequencing using Guppy v4.2.2 with the ‘--fast5_out’ option to retain raw signal data for downstream analyses. FASTQ reads were demultiplexed into biological replicates using Deeplexicon (Smith et al., 2020). We note that basecalling accuracy is largely irrelevant for signal-based methods.

### Read Alignment and Filtering

We aligned reads to *E.coli* K-12 MG1655 16S (*rrsE*, operon *rrnE*) and 23S (*rrlB*, operon *rrnB*) rRNA reference sequences using minimap2 v2.24 (H. Li, 2018) with the parameters ‘-ax splice -uf -k14 -- secondary=no’. Reference coordinates for 16S and 23S rRNA include a 200-nucleotide padding upstream of the annotated gene start and downstream of annotated gene end, to accommodate aligned reads extending beyond the reference boundaries. For all downstream analysis calculations only coordinates pertaining to the rRNA were used.

Aligned reads were processed using SAMtools v.1.17 (Danecek et al., 2021; H. Li et al., 2009).

For estimating the effect of coverage on different RNA modification detection tools, we subsampled reads to different target coverages (5×, 10×, 15×, 20×, 25×, 30×, 35×, 40×, 45×, 50×, 55×, 60×, 65×, 70×, 75×, 80×, 85×, 90×, 95×, 100×, 150×, 200×, 500×, 750×, 1000×) per rRNA reference. In addition to these subsampled levels, the full dataset without subsampling was also analysed. We calculated the target number of bases as (rRNA length x target depth) and subsampled an equivalent number of bases from the aligned reads using Filtlong v0.2.1 (Wick, 2017/2022) with options --min_length 500 --length_weight 10 --mean_q_weight 1 --window_q_weight 1 applied independently per replicate.

### Raw signal preprocessing

We converted multi-read FAST5 files to single-read FAST5 format using the ONT-Fast5-API v4.1.0 ‘multi_to_single_fast5ʹ module, and the reads mapping to 16S and 23S rRNA references were extracted using ‘fast5_subset’. Mapped read ids were extracted from BAM files using Seqkit v.2.3.1 (Shen et al., 2016). For signal-level analyses which require event-to-reference sequence alignment, we used f5c v1.5; (Gamaarachchi et al., 2020) to index FAST5 and FASTQ files (f5c index) and to generate signal-level event alignments (f5c eventalign) with the parameters ‘--rna --scale-events --signal-index --print-read-names --samples’. We performed a separate ‘f5c evenalign’ run with parameters ‘--rna --scale-events --signal-index’ (without --print-read-names) to produce output compatible with the tool xPore.

### RNA modification detection

We evaluated ten tools for their ability to detect RNA modifications by comparing native (modified) direct RNA sequencing data with *in vitro* transcribed (IVT; unmodified) control data. These tools broadly fall into two categories based on their detection strategy: (i) signal-level approaches that analyse raw nanopore current signals, and (ii) error-rate-based approaches that exploit systematic basecalling errors introduced by the presence of RNA modifications.

### Signal-level modification detection

**Tombo** v.1.5.1 (Stoiber et al., 2017). We ran Tombo using the ‘level_sample_compare’ method, which uses a two-sample Kolmogrov-Smirnov (KS) test at each reference position comparing the distribution of normalised current levels between native and IVT samples.

Raw signal data was re-squiggled to the reference sequences using ‘tombo resquiggle’ with the parameters ‘--rna --overwrite --num-most-common-errors 5’. We performed modification detection using ‘tombo detect_modifications level_sample_compare’ with ‘--statistic-type ks - -minimum-test-reads 1 --store-p-value’. Per-position statistics were then exported using Tombo Python API.

**Nanocompore** v1.0.4 (Leger et al., 2021). Eventalign data from f5c was first collapsed to per-position summary statistics using ‘nanocompore evenalign-collapse’. We then performed differential modification detection using ‘nanocompore sampcomp’ with parameters logistic regression (--logit), a sequence context of 2 bases (sequence_context 2), and a minimum coverage of 1 read per position (-- min_coverage 1).

**xPore** v2.1 (Pratanwanich et al., 2021). xPore uses a Bayesian Gaussian mixture model (GMM) for RNA modification detection. Eventalign data from f5c was first preprocessed using ‘xpore dataprep’, and then modification detection was done using ‘xpore diffmod’.

**Yanocomp** v0.2 (Parker et al., 2021a). Yanocomp uses a GMM-based approach with a KS test to identify modified positions. Eventalign data from f5c was converted to HDF5 format using ‘yanocomp prepare’, and modification detection was done using ‘yanocomp analysis’.

**nanoDoc** (Ueda, 2021). nanoDoc uses deep one-class classification with a convolution neural network (CNN) which analyses current signal deviations in native and IVT samples to give scores per-position. Initial data processing was done using ‘python nanoDoc.py formatfile’ and comparative analysis using ‘python nanoDoc.py analysis’ with additional parameters ‘-w’ (k-mer weights) and ‘-p’ (parameter set). Note that nanoDoc requires TensorFlow 2.3 with CUDA 10.1, which is generally incompatible with newer GPU architectures. Our initial runs were performed on a compatible GPU (NVIDIA GTX 1080Ti), but this hardware was no longer available for subsequent analyses. Our attempts to port the environment to newer GPUs were unsuccessful due to unresolvable dependency conflicts, so complete evaluation across the full parameter space was not possible.

### Error-rate-based modification detection

**ELIGOS2** v2.1.0 (Jenjaroenpun et al., 2021). We ran ELIGOS2 using ‘eligos2 pair_diff_mod’ to compare per-position basecalling error patterns between native and IVT BAM files with ‘--min_depth 1’.

**EpiNano** v1.2.5 (Liu et al., 2019). EpiNano has two modes - SVM and Error. We used the Error mode throughout the analysis as this is the mode which is modification agnostic. Per-position error metrics (mismatch, insertion, and deletion frequencies) were extracted from native and IVT BAM files using ‘Epinano_Variants.py’. We then performed differential error analysis using ‘Epinano_DiffErr.R’ in two modes: mismatch-only (-f mis) and sum-of-errors (-f sum_err, using preprocessed data from ‘Epinano_sumErr.py’), with an error rate difference threshold of 0.1 (-d 0.1). We used sum-of-errors mode outputs throughout the main analyses.

**DiffErr** (Parker et al., 2020). We ran DiffErr to compare read-level error patterns between conditions, outputting per-position odds ratios, G-statistics, and p-values, with additional parameters ‘--median-expr-threshold 0’, ‘--min-expr-threshold 0’, and ‘-f 1.0’ (FDR threshold set to 1.0 to retain all tested positions). DiffErr reports both a G-statistic and a -log10(FDR) for each position. We used the G-statistic as the scoring metric because at moderate-to-high coverage (>30x), nearly all positions reached extreme significance (-log10(FDR) > 300), producing identical scores at modified and unmodified sites. This made it difficult to rank modified above unmodified sites, whereas the G-statistic retains discriminative power across coverage levels.

**DRUMMER** (Abebe et al., 2022). We ran DRUMMER to compare per-position base fraction distributions between native and IVT samples using a G-test, with ‘-a exome’ (appropriate for targeted reference analysis).

**JACUSA2** v2.0.4; (Piechotta et al., 2022). JACUSA2 uses a pileup-based likelihood scoring approach to detect positions with significant differences in base composition between conditions. We ran the tool in pairwise comparison mode (call-2) with parameters ‘-c 1’ (minimum coverage), ‘-m 0’ (minimum score of 0), and ‘-q 0’ (minimum base quality of 0).

### Benchmarking Parameter Configuration

For unbiased benchmarking across all tools and to construct receiver operating characteristic (ROC) and precision-recall (PR) curves across full range of score thresholds, we set all tool-specific filtering thresholds to maximally permissive values so that all tested positions in ribosomal RNAs were reported in each tool’s output. Specifically, p-value and false discovery rate (FDR) thresholds were set to 1.0, minimum coverage thresholds were set to 0 or 1, and odds ratio thresholds were set to 0 for tools that have these options configurable. This approach ensured that performance evaluation reflects the intrinsic discriminative ability of each tool’s output statistic rather than the choice of arbitrary cutoffs, consistent with the strategies employed by (Maestri et al., 2024) and (Zhong et al., 2023a).

### Downstream Benchmarking Analysis

We used custom Python scripts (https://github.com/bhargava-morampalli/rnamod-tool-benchmark-code) to standardise, compare, and benchmark outputs across tools. All analyses were performed using Python (≥3.10), pandas (≥1.5.0), NumPy (≥1.24.0), scikit-learn (≥1.3.0), matplotlib (≥3.7.0), and seaborn (≥0.12.0), and SciPy (≥1.10.0).

### Output parsing and standardisation

We parsed tool outputs into a common schema containing: tool name, reference (16S and 23S), reference position (1-based), primary score, score type (p-value, FDR, z-score, -log10 FDR, or direct score), and replicate identifier.

For each tool and replicate, parsed outputs were standardised to the full evaluation region of each rRNA (1542 positions for 16S and 2904 positions for 23S). To quantify tool output completeness, we calculated the overall call fraction (OCF) for each tool as the proportion of positions within the evaluation region for which any score was reported. We evaluated metrics under two scoping strategies: 1) ‘all-position’ scope, where unreported positions were imputed with a score of zero before computing AUROC and AUPRC, and 2) ‘reported-position-only’ scope, where metrics computed only over positions that the tool reported. The difference between these two approaches quantifies the extent to which incomplete output inflates apparent performance.

### Score Directionality and Transformation

As the tools output different statistical scores, we transformed scores so that larger values consistently indicate modified positions.

| Tool | Output statistic | Transformation |
| --- | --- | --- |
| Tombo | p-value | $-\log_{10}(\text{p-value})$ |
| Nanocompore | GMM logit p-value | $-\log_{10}(\text{p-value})$ |
| xPore | p-value | $-\log_{10}(\text{p-value})$ |
| nanoDoc | Raw distance score | Used directly |
| Yanocomp | FDR | $-\log_{10}(\text{FDR})$ |
| ELIGOS2 | Adjusted p-value | $-\log_{10}(\text{adj p-value})$ |
| EpiNano | z-score | $\text{abs}(\text{z-score})$ |
| DiffErr | $-\log_{10}(\text{FDR})$ | Used directly |
| DiffErr (G-stat) | G-statistic | Used directly |
| DRUMMER | Adjusted p-value (G-test) | $-\log_{10}(\text{adj p-value})$ |
| JACUSA2 | Pileup score | Used directly |

The G-statistic was used as the primary DiffErr score for all main analyses; see DiffErr dual-score evaluation below.

### Ground Truth Definition

We compiled known modification sites in *E. coli* 16S and 23S rRNA from the MODOMICS database (Boccaletto et al., 2022). The ground truth comprised 11 modified positions in 16S rRNA and 25 modified positions in 23S rRNA, encompassing diverse modification types (**Supp. Tables. S1** and **S2**). We classified each position within the evaluation region (see Position Standardisation above) as either modified (positive) or unmodified (negative) based on the ground truth, a binary classification task with 16S rRNA: 11 positives, 1,531 negatives (1,542 total positions) and 23S rRNA: 25 positives, 2,879 negatives (2,904 total positions).

## Benchmark Metrics and Statistical Analysis

### Performance Metrics

We evaluated tool performance using area under the receiver operating characteristic curve (AUROC) and area under the precision-recall curve (AUPRC) using scikit-learn’s ‘roc_auc_score’ and ‘average_precision_score’ functions, respectively. Precision, recall, and F1 score were computed at a threshold that maximised the F1 score on the precision-recall curve. The maximal F1 threshold was determined by computing F1 across all unique score values for each tool and selecting the threshold that yielded the maximum F1, with:

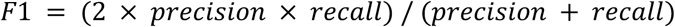

Or alternatively,

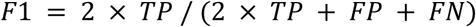

### ROC Curve Analysis

We generated ROC curves to evaluate the trade-off between sensitivity (true positive rate = TP/(TP + FN) and specificity = TN/(TN+FP)) for each tool. Each tool’s transformed output statistic (see Score Directionality table above) served as the classification threshold.

### Precision-Recall Curve Analysis

Due to strong class imbalance in the dataset (few modification positions vs many unmodified positions), we also generated precision-recall (PR) curves and calculated the Area Under the Precision-Recall Curve (AUPRC) alongside ROC curves. Under severe class imbalance, AUROC can appear inflated by the large number of true negatives, whereas AUPRC directly reflects performance of the minority (modified) class.

### Replicate Handling

We computed AUROC and AUPRC separately for each biological replicate and then averaged them. Median, standard deviation, and per-replicate values were also calculated to show variability.

### Window-Size Analysis

To account for potential positional offsets in modification detection (due to the 5 nucleotides occupying the pore simultaneously in nanopore sequencing), we evaluated precision using window sizes of 0 (exact match) to 10 nucleotides. For a given window size of n, each ground-truth modification position was expanded to include all positions within ± n positions. A tool prediction falling within this expanded set was counted as true positive. This approach is consistent with the nucleotide tolerance windows used in comparable studies (Jenjaroenpun et al., 2021; Guo et al., 2025; Spangenberg et al., 2025). We recomputed AUROC, AUPRC, and F1 using window sizes of 0 (exact match) to 10 nucleotides.

### Position Offset Analysis

To detect systematic positional biases in tool predictions, we performed a lag scan by uniformly shifting all ground-truth modification positions by δ = -10 to +10 nucleotides and recomputing AUROC and AUPRC at each offset. Unlike symmetric window tolerance analysis, the lag scan applied a fixed directional shift, enabling detection of tools whose modification predictions are systematically displaced relative to annotated modification sites. To evaluate whether systematic positional offsets in tool predictions could be corrected to improve site-level recovery, we performed an offset-corrected detection analysis at 1000× coverage. For each tool, the peak offset δ was defined as the shift (from the lag scan analysis described above) that maximised mean AUPRC across replicates, independently for each rRNA molecule. For tools where δ ≠ 0, we evaluated detection at the shifted position (ground-truth positions + δ) using the maximum F1 threshold determined at that offset from the lag analysis. A site was classified as detected if its score at the shifted position exceeded the offset-specific threshold. Consensus detection required detection in ≥ 2 replicates (majority rule). Results were compared against standard exact-position (δ=0) evaluation to quantify sites gained and lost.

### Tool Combination Analysis

To identify optimal tool combinations for modification site recovery, we computed the union of positive calls across all possible single tools, pairs (45 combinations), and triples (120 combinations) at 1000× coverage. A position was classified as a true positive if it corresponded to a ground-truth site accounting for each tool’s offset (at δ=0 for exact-match evaluation, or at the tool’s peak offset for offset-corrected evaluation). False positives were positions called positive by any tool in the combination that did not correspond to any ground-truth site. True positive and false positive counts were averaged across three biological replicates. The Pareto frontier was computed as the set of combinations for which no alternative achieved both higher site recovery and lower false positives.

## Data and Code Availability

Raw FAST5 files containing the raw signal data used have been deposited in the European Nucleotide Archive (ENA) under study accession PRJEB58726. Processed benchmarking outputs, including per-tool score files, aggregated performance metrics, and all supplementary tables, are available at https://github.com/bhargava-morampalli/rnamod-tool-benchmark-code.

The Nextflow pipeline used to process raw sequencing data from mapping to modification detection and downstream metric computation is available at https://github.com/bhargava-morampalli/rnamodbench. Python scripts for all visualisations, statistical analyses, offset-corrected site-level recovery analysis, and tool combination analysis are available at https://github.com/bhargava-morampalli/rnamod-tool-benchmark-code.

## Acknowledgements

We thank N. Freed for input into the protocol for producing unmodified RNA.

## Author Notes

OKS has received funding from ONT for travel and accommodation.

BRM and OKS conceived and designed the experiments. BRM performed the experiments. BRM performed the analyses with input from OKS. BRM and OKS wrote the manuscript.

## Supplementary Tables

**Supplementary Table S1.** Known modifications on *Escherichia coli* 16S ribosomal RNA. The first column indicates the position relative to the transcription start. The second column indicates the specific type of modification. The majority of modifications are toward the 3ʹ end of the molecule.

| Position | Modification |
| --- | --- |
| 516 | Ψ |
| 527 | m<sup>7</sup>G |
| 966 | m<sup>2</sup>G |
| 967 | m<sup>5</sup>C |
| 1207 | m<sup>2</sup>G |
| 1402 | m<sup>4</sup>Cm |
| 1407 | m<sup>5</sup>C |
| 1498 | m<sup>3</sup>U |
| 1516 | m<sup>2</sup>G |
| 1518 | m<sup>6</sup><sub>2</sub>A |
| 1519 | m<sup>6</sup><sub>2</sub>A |

**Supplementary Table S2.** Known modifications on *Escherichia coli* 23S ribosomal RNA. The first column indicates the position relative to the transcription start. The second column indicates the specific type of modification. As with 16S, the majority of modifications are toward the 3ʹ end of the molecule.

| Position | Modification |
| --- | --- |
| 745 | m<sup>1</sup>G |
| 746 | ψ |
| 747 | m<sup>5</sup>U |
| 955 | ψ |
| 1618 | m<sup>6</sup>A |
| 1835 | m<sup>2</sup>G |
| 1911 | ψ |
| 1915 | m<sup>3</sup>ψ |
| 1917 | ψ |
| 1939 | m<sup>5</sup>U |
| 1962 | m<sup>5</sup>C |
| 2030 | m<sup>6</sup>A |
| 2069 | m<sup>7</sup>G |
| 2251 | Gm |
| 2445 | m<sup>2</sup>G |
| 2449 | hU |
| 2457 | ψ |
| 2498 | Cm |
| 2501 | oh<sup>5</sup>C |
| 2503 | m<sup>2</sup>A |
| 2504 | ψ |
| 2552 | Um |
| 2580 | ψ |
| 2604 | ψ |
| 2605 | ψ |

**Supplementary Table S3:**
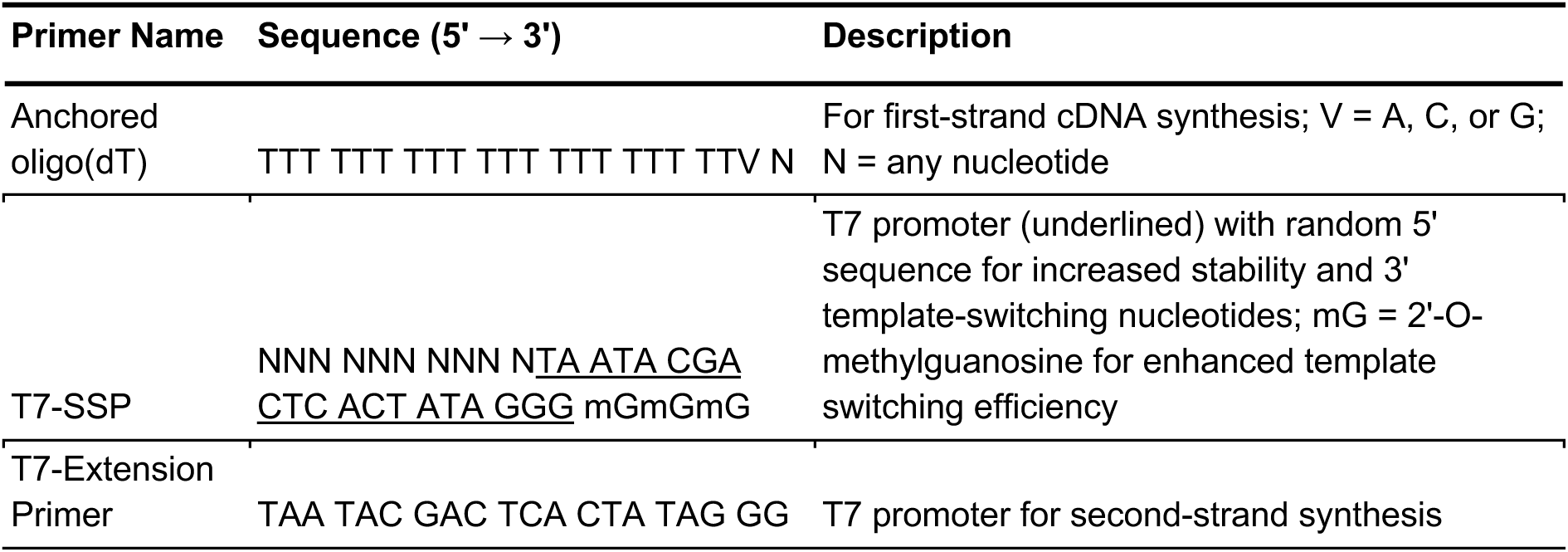
Custom oligos for modified strand switch protocol for unmodified RNA synthesis.

**Supplementary Table S4:** AUROC performance of RNA modification detection tools across coverage levels. Minimum coverage at which mean AUROC first exceeds 0.9 across subsampled replicates (5×-1000×). “>1000×” indicates the tool did not reach 0.9 within the subsampled range. Full coverage corresponds to the complete, subsampled dataset. nanoDoc was not tested at full coverage due to environment incompatibilities (see text). Tools are ordered by the earliest threshold reached.

| Tool | Min. Coverage for AUROC > 0.9 |  | AUROC at 1000x |  | AUROC at full coverage |  |
| --- | --- | --- | --- | --- | --- | --- |
|  | 16S rRNA | 23S rRNA | 16S rRNA | 23S rRNA | 16S rRNA | 23S rRNA |
| DiffErr | 25× | 30× | 0.944 | 0.955 | 0.966 | 0.960 |
| JACUSA2 | 45× | 75× | 0.910 | 0.955 | 0.990 | 0.957 |
| ELIGOS2 | 90× | 45× | 0.869 | 0.926 | 0.889 | 0.931 |
| DRUMMER | >1000× | >1000× | 0.742 | 0.731 | 0.803 | 0.658 |
| Tombo | >1000× | 750× | 0.867 | 0.899 | 0.877 | 0.896 |
| nanoDoc | >1000× | >1000× | 0.799 | 0.830 | N/A | N/A |
| Nanocompore | >1000× | >1000× | 0.634 | 0.762 | 0.853 | 0.775 |
| EpiNano | >1000× | >1000× | 0.663 | 0.877 | 0.765 | 0.901 |
| Yanocomp | >1000× | >1000× | 0.671 | 0.836 | 0.958 | 0.855 |
| xPore | >1000× | >1000× | 0.660 | 0.623 | 0.637 | 0.535 |

**Supplementary Table S5.** Effect of offset correction on tool performance at 1000× coverage for 16S. For each tool, performance metrics are shown at exact-position evaluation (δ = 0) and at each tool’s empirically determined peak offset (Peak Offset). Metrics include AUPRC, Precision, Recall, F1 score at the maximal F1 threshold, as well as ground-truth sites recovered (TP) and false positives (FP). Tools with peak offset of 0 (DiffErr, JACUSA2, DRUMMER) show identical values in both columns. TP and FP are mean values across three biological replicates.

| Tool | Ground Truth Sites (n) | Peak Offset ($\delta$) | AUPRC | | Precision | | Recall | | F1 | | Sites Recovered | | False Positives | |
| --- | --- | --- | --- | --- | --- | --- | --- | --- | --- | --- | --- | --- | --- | --- |
| | | | $\delta = 0$ | Peak Offset | $\delta = 0$ | Peak Offset | $\delta = 0$ | Peak Offset | $\delta = 0$ | Peak Offset | $\delta = 0$ | Peak Offset | $\delta = 0$ | Peak Offset |
| DiffErr | 11 | 0 | 0.536 | 0.536 | 0.769 | 0.769 | 0.606 | 0.606 | 0.677 | 0.677 | 6.7 | 6.7 | 2 | 2 |
| JACUSA2 | 11 | 0 | 0.478 | 0.478 | 0.889 | 0.889 | 0.455 | 0.455 | 0.6 | 0.6 | 5 | 5 | 0.6 | 0.6 |
| DRUMMER | 11 | 0 | 0.379 | 0.379 | 0.708 | 0.708 | 0.515 | 0.515 | 0.596 | 0.596 | 5.7 | 5.7 | 2.3 | 2.3 |
| ELIGOS2 | 11 | -1 | 0.103 | 0.164 | 0.187 | 0.75 | 0.333 | 0.182 | 0.234 | 0.245 | 3.7 | 2 | 16 | 0.7 |
| EpiNano | 11 | -1 | 0.02 | 0.408 | 0.089 | 0.49 | 0.091 | 0.545 | 0.09 | 0.515 | 1 | 6 | 10.2 | 6.3 |
| Tombo | 11 | -4 | 0.052 | 0.185 | 0.068 | 0.237 | 0.606 | 0.545 | 0.122 | 0.293 | 6.7 | 6 | 91.7 | 19.4 |
| Nanocompore | 11 | -2 | 0.057 | 0.219 | 0.192 | 0.792 | 0.273 | 0.212 | 0.22 | 0.31 | 3 | 2.3 | 12.6 | 0.6 |
| nanoDoc | 11 | -4 | 0.049 | 0.273 | 0.082 | 0.255 | 0.364 | 0.667 | 0.132 | 0.368 | 4 | 7.3 | 44.5 | 21.4 |
| xPore | 11 | -1 | 0.099 | 0.107 | 0.369 | 0.467 | 0.212 | 0.182 | 0.267 | 0.257 | 2.3 | 2 | 4 | 2.3 |
| Yanocomp | 11 | -1 | 0.08 | 0.138 | 0.355 | 0.288 | 0.182 | 0.333 | 0.213 | 0.306 | 2 | 3.7 | 3.6 | 9.1 |

**Supplementary Table S6.** Effect of offset correction on tool performance at 1000× coverage for 23S. For each tool, performance metrics are shown at exact-position evaluation (δ = 0) and at each tool’s empirically determined peak offset (Peak Offset). Metrics include AUPRC, Precision, Recall, F1 score at the maximal F1 threshold, as well as ground-truth sites recovered (TP) and false positives (FP). Tools with peak offset of 0 (DiffErr, JACUSA2, DRUMMER) show identical values in both columns. TP and FP are mean values across three biological replicates.

| Tool | Ground Truth Sites (n) | Peak Offset ($\delta$) | AUPRC | | Precision | | Recall | | F1 | | Sites Recovered | | False Positives | |
| --- | --- | --- | --- | --- | --- | --- | --- | --- | --- | --- | --- | --- | --- | --- |
| | | | $\delta = 0$ | Peak Offset | $\delta = 0$ | Peak Offset | $\delta = 0$ | Peak Offset | $\delta = 0$ | Peak Offset | $\delta = 0$ | Peak Offset | $\delta = 0$ | Peak Offset |
| DiffErr | 25 | 0 | 0.46 | 0.46 | 0.385 | 0.385 | 0.64 | 0.64 | 0.48 | 0.48 | 16 | 16 | 25.6 | 25.6 |
| JACUSA2 | 25 | 0 | 0.52 | 0.52 | 0.48 | 0.48 | 0.587 | 0.587 | 0.526 | 0.526 | 14.7 | 14.7 | 15.9 | 15.9 |
| DRUMMER | 25 | 0 | 0.443 | 0.443 | 0.814 | 0.814 | 0.453 | 0.453 | 0.581 | 0.581 | 11.3 | 11.3 | 2.6 | 2.6 |
| ELIGOS2 | 25 | 0 | 0.268 | 0.268 | 0.29 | 0.29 | 0.587 | 0.587 | 0.387 | 0.387 | 14.7 | 14.7 | 36 | 36 |
| EpiNano | 25 | -1 | 0.173 | 0.505 | 0.284 | 0.701 | 0.4 | 0.467 | 0.331 | 0.56 | 10 | 11.7 | 25.2 | 5 |
| Tombo | 25 | -2 | 0.107 | 0.151 | 0.196 | 0.269 | 0.293 | 0.413 | 0.233 | 0.325 | 7.3 | 10.3 | 30 | 28.1 |
| Nanocompore | 25 | -1 | 0.124 | 0.218 | 0.32 | 0.278 | 0.267 | 0.427 | 0.288 | 0.287 | 6.7 | 10.7 | 14.2 | 27.7 |
| nanoDoc | 25 | -4 | 0.072 | 0.237 | 0.137 | 0.236 | 0.44 | 0.507 | 0.208 | 0.318 | 11 | 12.7 | 69.4 | 40.9 |
| xPore | 25 | 1 | 0.073 | 0.109 | 0.226 | 0.211 | 0.28 | 0.387 | 0.25 | 0.273 | 7 | 9.7 | 23.9 | 36.1 |
| Yanocomp | 25 | -1 | 0.108 | 0.128 | 0.184 | 0.209 | 0.4 | 0.4 | 0.247 | 0.274 | 10 | 10 | 44.4 | 37.8 |

## Supplementary Figures

**Supplementary Figure S1.**
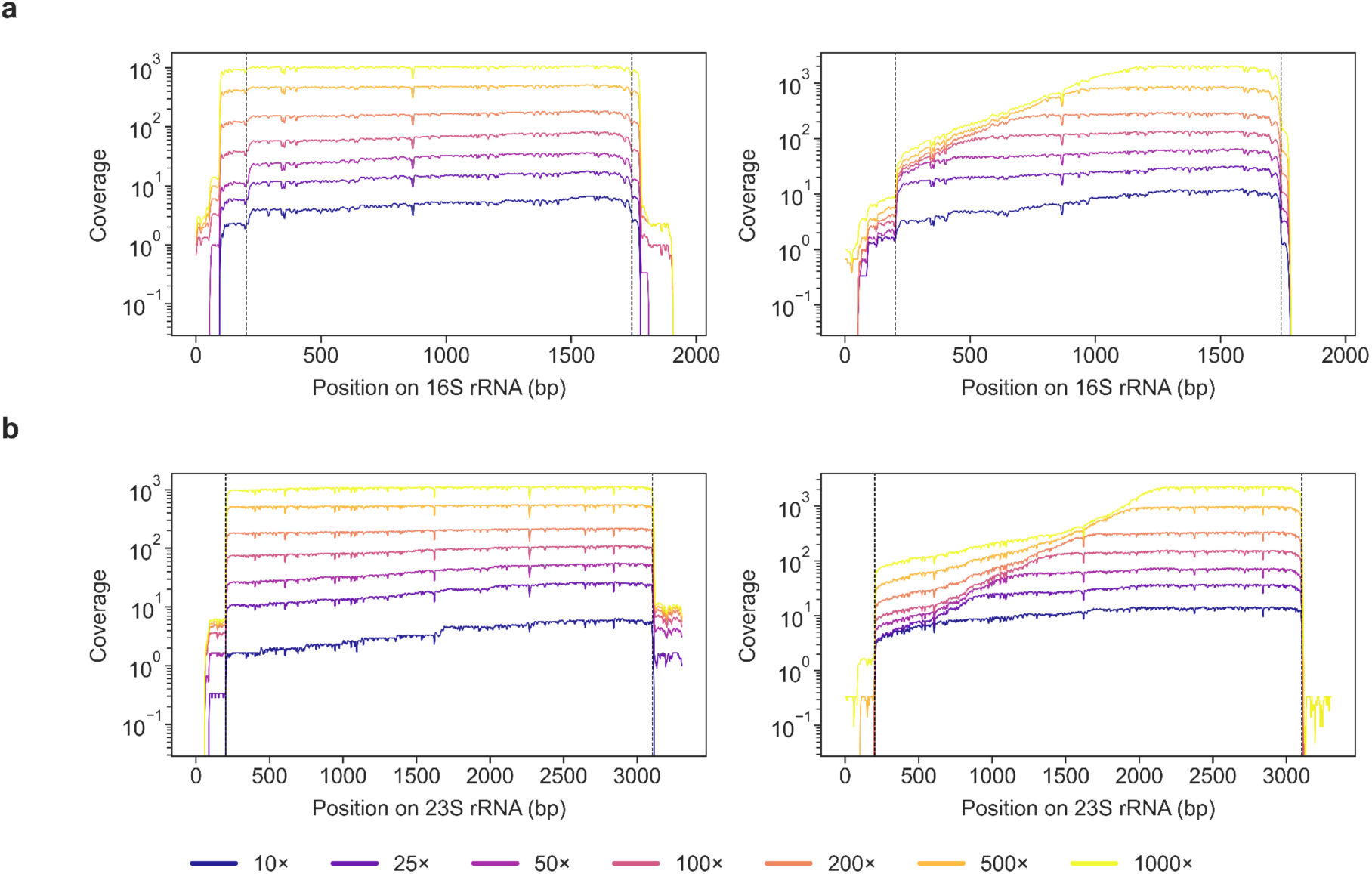
Coverage uniformity improves with subsampling. Coverage across (**a)** native 16S (left) and IVT 16S (right), (**b)** native 23S (left) and IVT 23S (right) rRNA at seven target coverage levels. Lines show mean coverage across three biological replicates, smoothed with a 7 bp window. Dashed vertical lines mark transcript boundaries. Native rRNA coverage is relatively uniform across all depths. IVT coverage still shows 3’ bias, reflecting truncation during reverse transcription rather than insufficient sequencing depth.

**Supplementary Figure S2.**
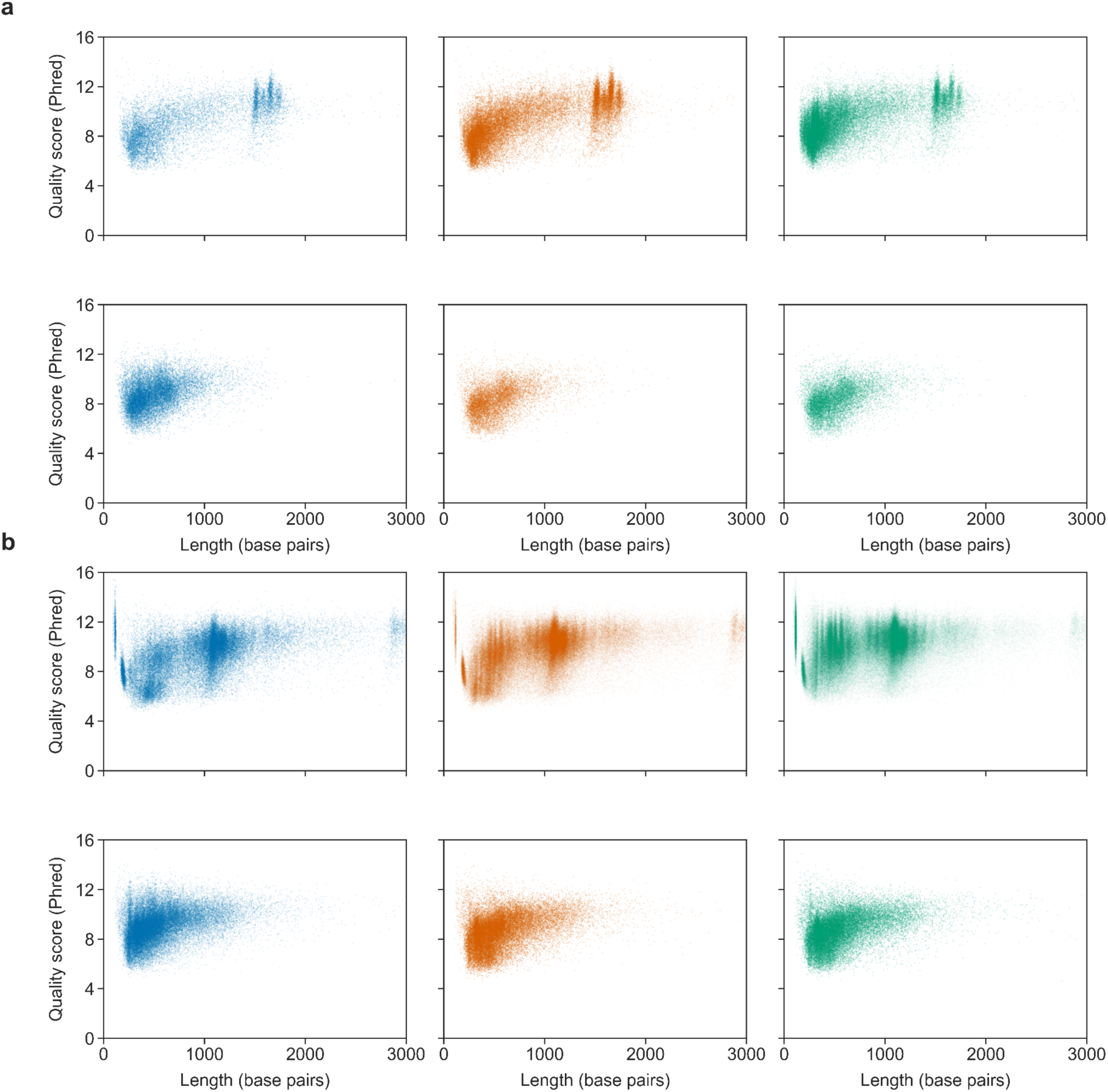
Read quality is comparable between native and IVT rRNA at equivalent read lengths. Read quality (Phred score) versus read length for (**a**) 16S and (**b)** 23S rRNA. For each panel, native (upper) and IVT (lower) reads are shown across three biological replicates (columns). Point opacity scales with read count to maintain visibility across panels with different sequencing depths.

**Supplementary Figure S3.**
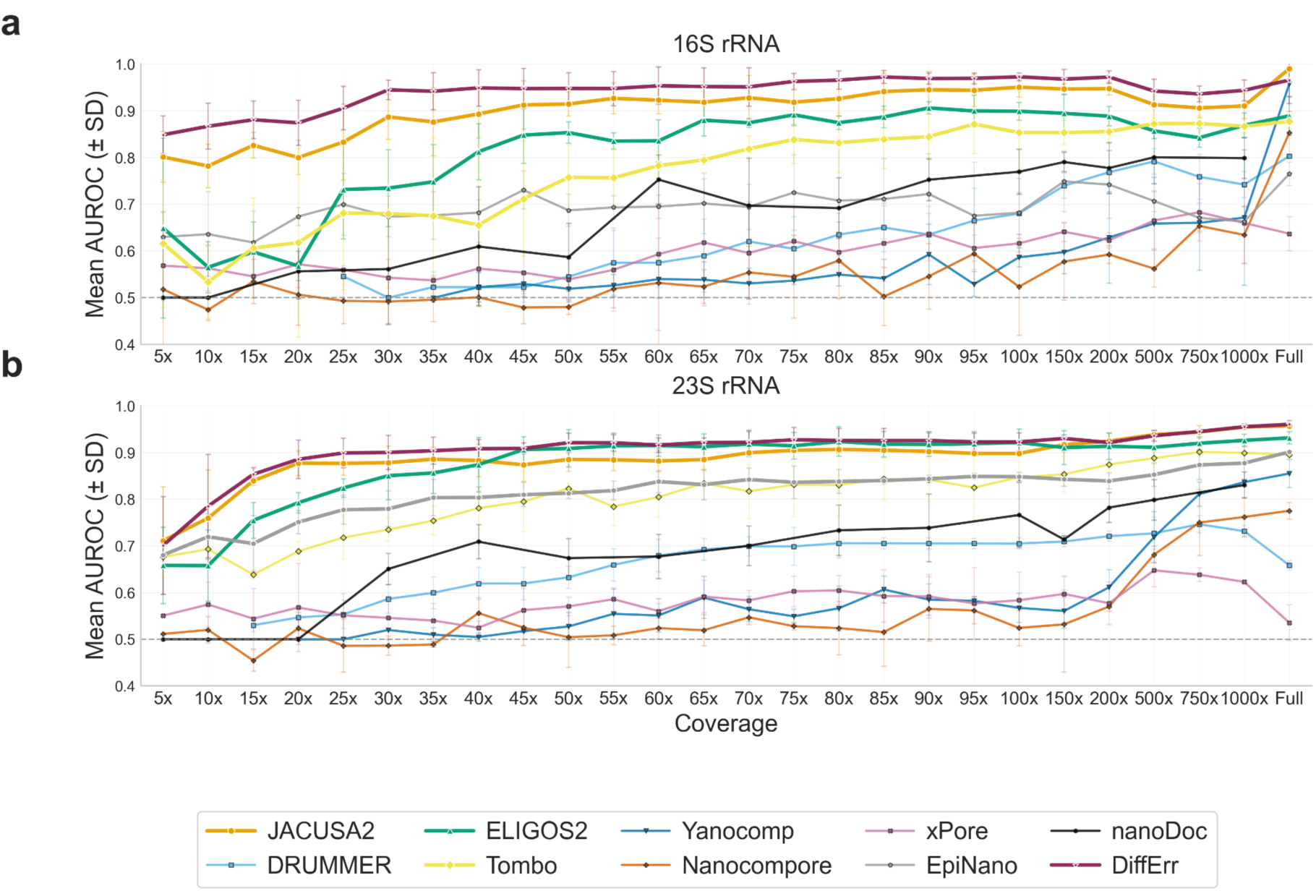
Coverage-dependent AUROC across all tools (overlay view). Mean AUROC for all nine tools plotted together for a) 16S rRNA b) 23S rRNA. The four highest ranked tools by mean AUROC are represented by thick lines; remaining tools are shown thinner lines. Error bars indicate ±SD; dashed line marks AUROC = 0.5.

**Supplementary Figure S4.**
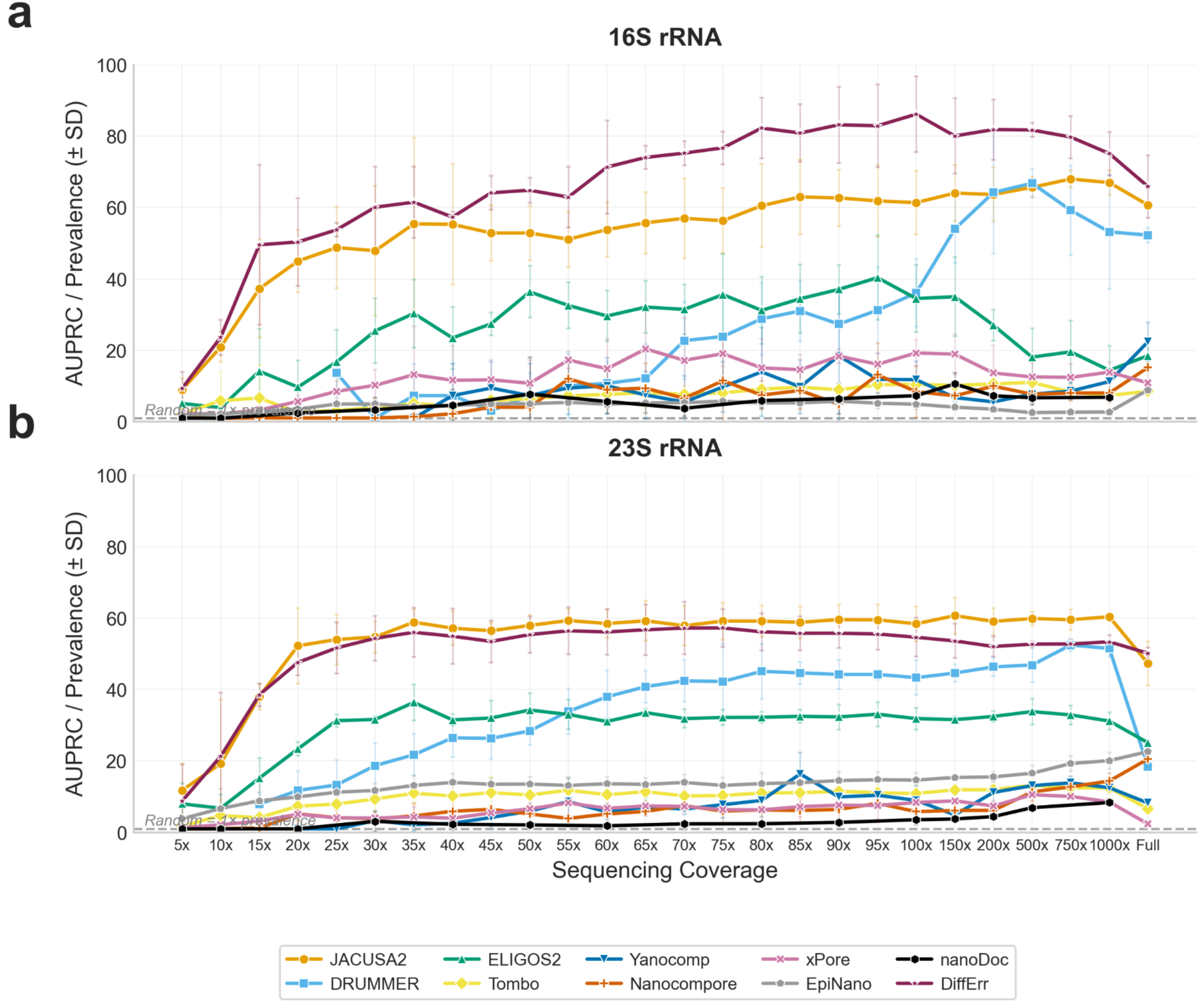
Fold improvement in AUPRC over random baseline across coverage levels. AUPRC normalised by modification prevalence (fold = AUPRC/prevalence) for all nine tools. Top panel (a) shows 16S rRNA and bottom panel (b) shows 23S rRNA. A fold value of 1 indicates random classifier performance. JACUSA2, DRUMMER and ELIGOS2 are top three performers, while most remaining tools remain low.

**Supplementary Figure S5.**
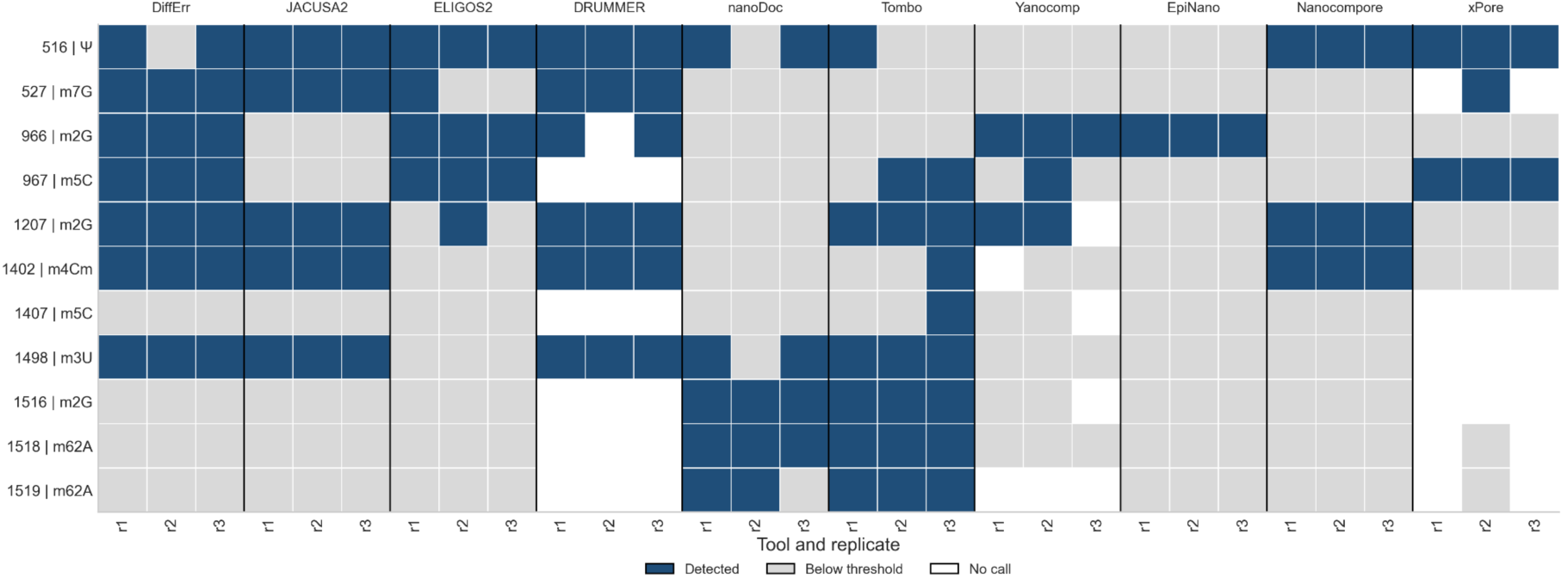
Each cell shows one replicate of a tool in which a known modification site was detected at that tool’s maximal F1 threshold for 16S rRNA. Blue indicates detected, grey indicates below the threshold and white indicates non detected. Sites are ordered by genomics position (rows) and each column indicates a replicate for that tool (indicated above each 3 replicate group). Site labels indicate position and modification type.

**Supplementary Figure S6.**
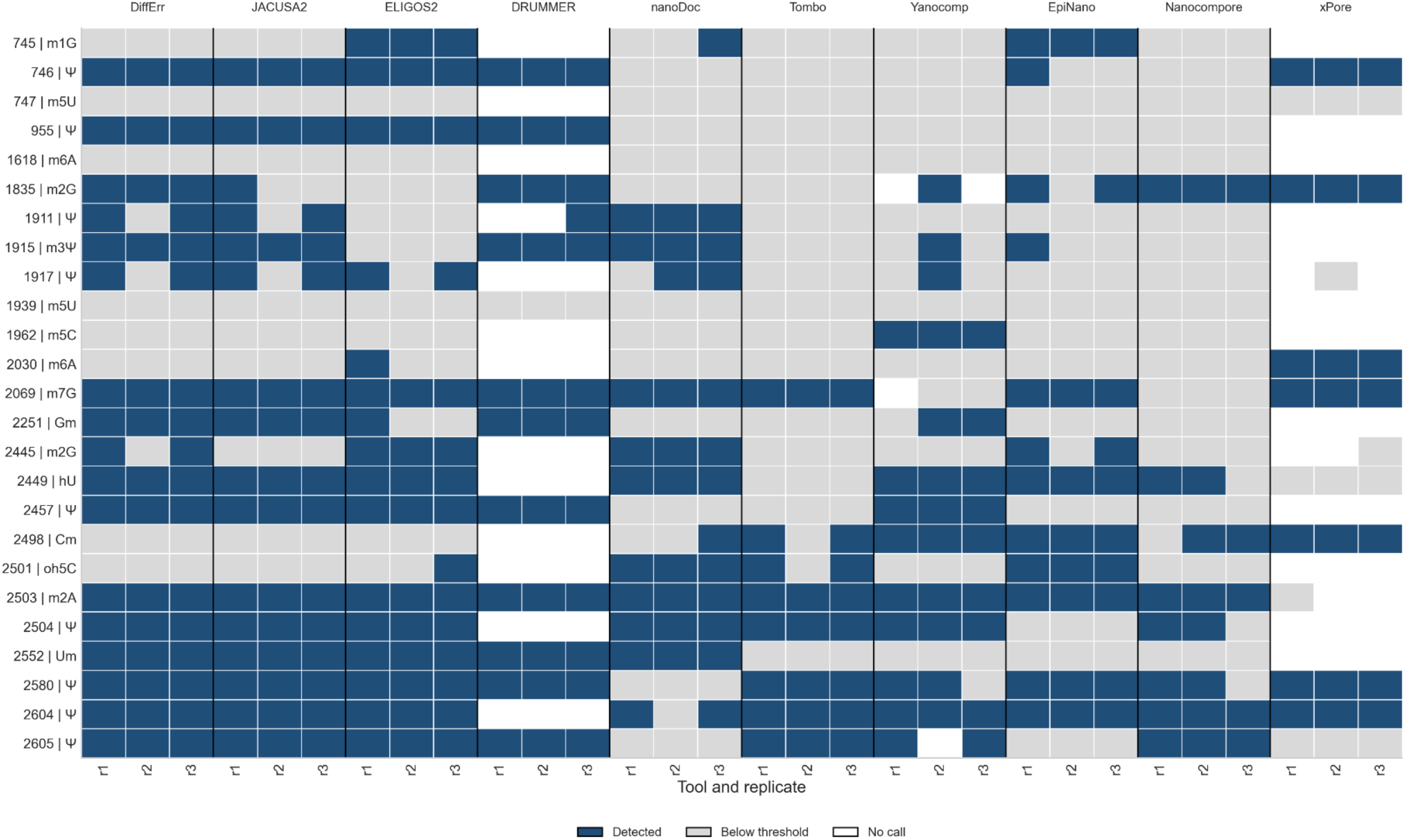
Each cell shows one replicate of a tool in which a known modification site was detected at that tool’s maximal F1 threshold for 23S rRNA. Blue indicates detected, grey indicates below the threshold and white indicates not detected. Sites are ordered by genomics position (rows) and each column indicates a replicate for that tool (indicated above each 3 replicate group). Site labels indicate position and modification type.

**Supplementary Figure S7.**
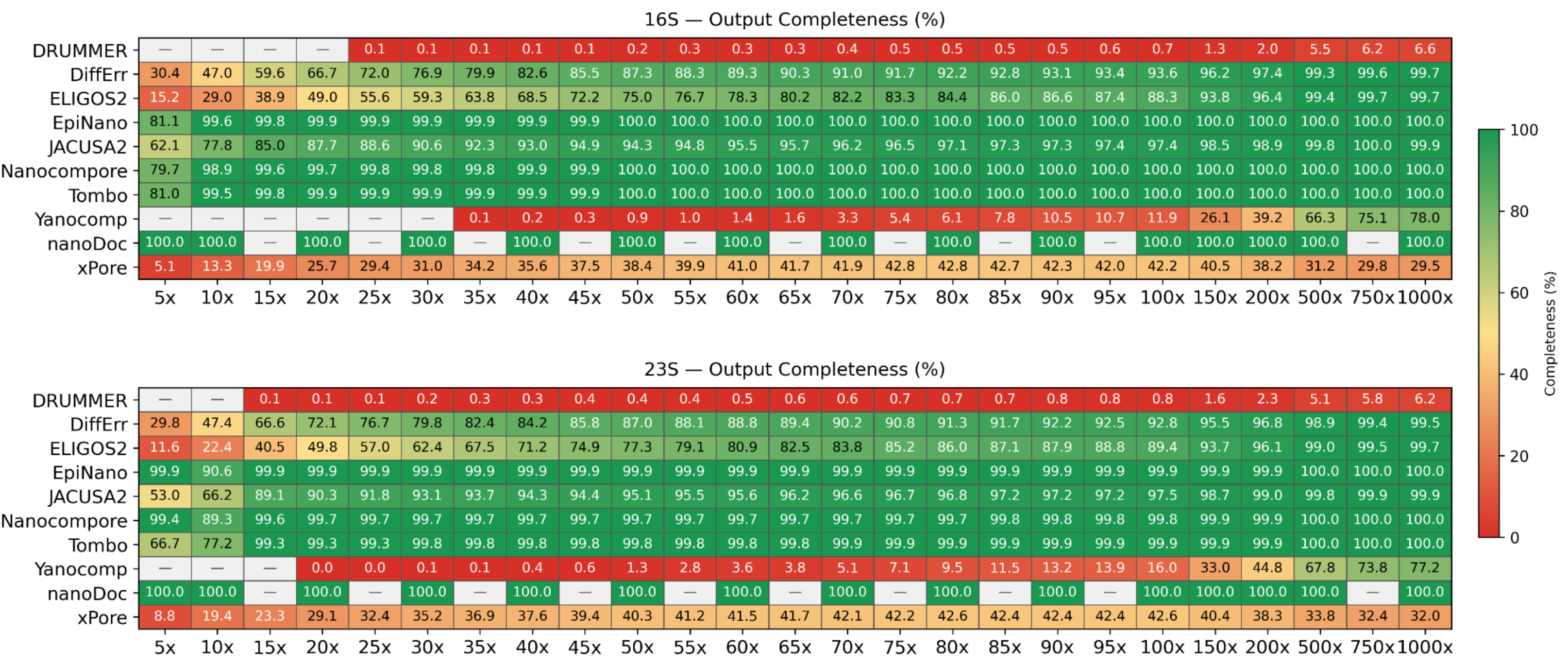
Output completeness of RNA modification detection tools at different coverages. Heatmaps show the percentage of evaluated positions for which each tool reports a score, across 25 coverage depths (5x-1000×) on *E. coli* 16S rRNA (top) and 23S rRNA (bottom). Colour scale ranges from red (0%) through yellow (50%) to green (100%). Grey cells with em-dash indicate coverages where a tool failed to produce output and in the case of the tool nanoDoc, it indicates coverages not tested

**Supplementary Figure S8.**
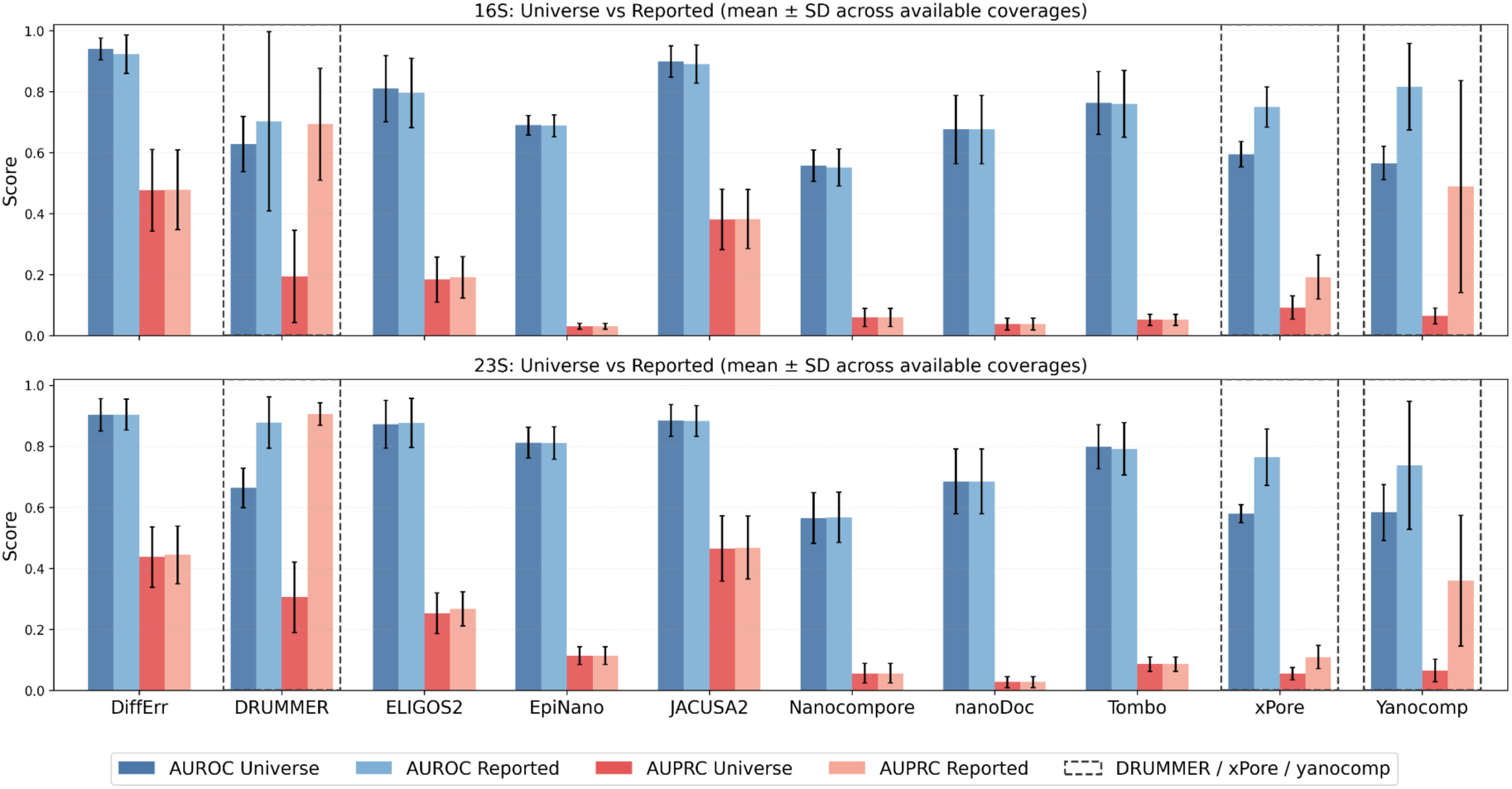
Discrimination performance of RNA modification detection tools across coverage levels. Mean AUROC and AUPRC (± SD) computed across 25 coverage depths (5x-1000×) for 10 tools benchmarked on *E. coli* 16S rRNA (top) and 23S rRNA (bottom). For each tool, four bars show AUROC and AUPRC evaluated over the fill universe of evaluated positions (all-position evaluation in darker shades) and over only positions the tool reported (Reported: lighter shades). Evaluable positions are defined as 16S rRNA,1,542 positions and 23S rRNA, 2,904 positions. Dashed boxes highlight DRUMMER, xPore, and Yanocomp, which exhibited large discrepancies between all-position and reported-position only discrimination metrics due to incompletely reported positions by these tools. Error bars represent standard deviation.

**Supplementary Figure S9.**
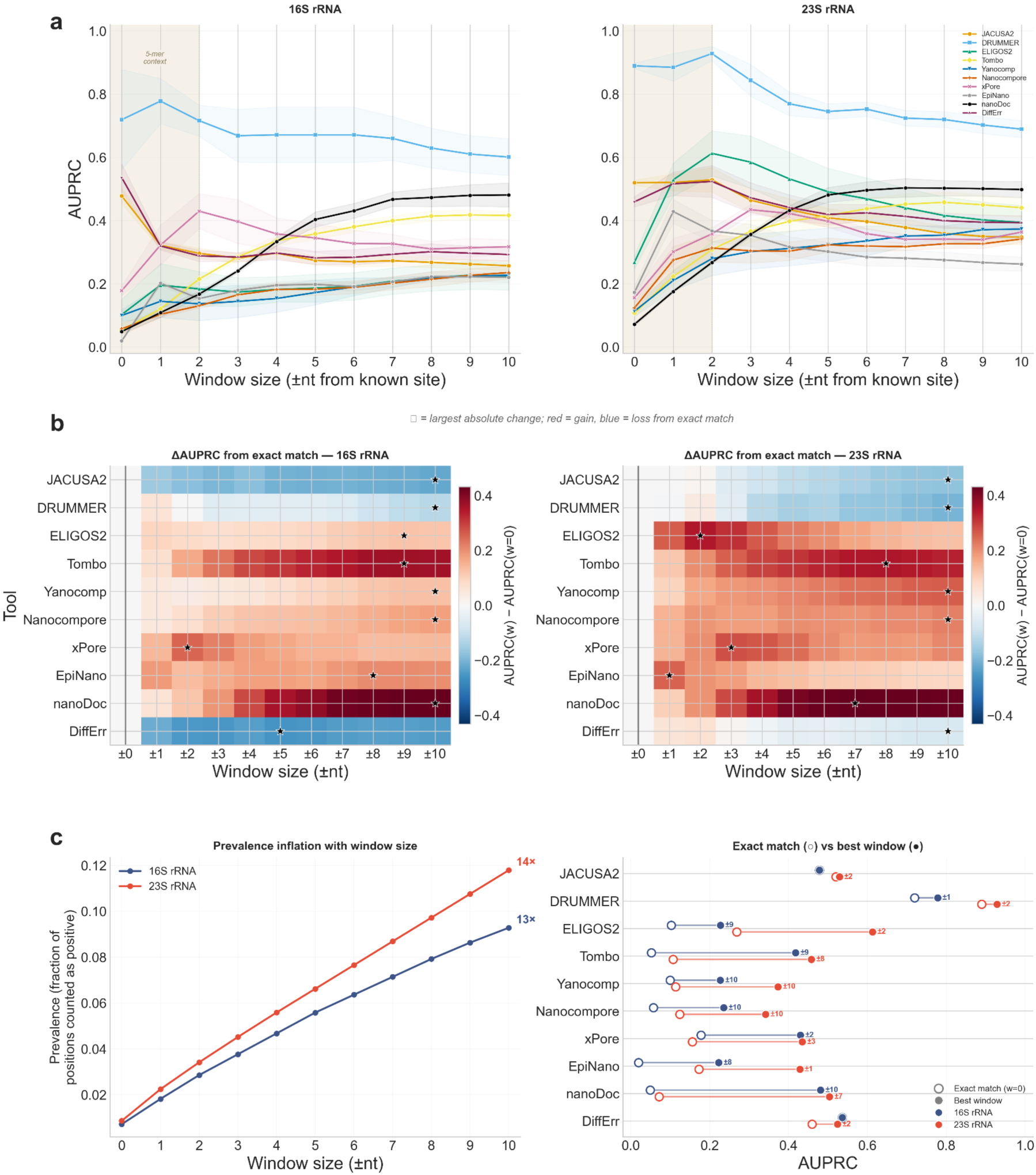
Symmetric tolerance windows inflate apparent performance and obscure positional precision differences between tools. Analysis performed at 1000× coverage on *E. coli* 16S rRNA (left) and 23S rRNA (right). **(a)** AUPRC as a function of symmetric tolerance window size (±0 to ±10 nt from known modification sites). Each line represents one tool; shaded bands indicate ±SD across biological replicates. The beige region indicates approximate 5-mer reader head context. **(b) δ** AUPRC heatmaps showing the change in AUPRC relative to exact-position evaluation (w = 0, vertical black line) for each tool and window size. Red indicates gains and blue indicates losses from the exact match. Stars mark the largest absolute change per tool. **(c, left)** Effective modification prevalence across window sizes, showing the fraction of positions counted as positive as windows expand. **(c, right)** Dumbbell chart comparing each tool’s AUPRC at exact position (open circles) with its peak AUPRC across all window sizes (filled circles), shown separately for 16S (blue) and 23S (red) rRNA.

## References

Abebe, J. S., Price, A. M., Hayer, K. E., Mohr, I., Weitzman, M. D., Wilson, A. C., & Depledge, D. P. (2022). DRUMMER—rapid detection of RNA modifications through comparative nanopore sequencing. Bioinformatics, 38(11), 3113–3115. 10.1093/bioinformatics/btac274

Anreiter, I., Mir, Q., Simpson, J. T., Janga, S. C., & Soller, M. (2021). New Twists in Detecting mRNA Modification Dynamics. Trends in Biotechnology, 39(1), 72–89. 10.1016/j.tibtech.2020.06.002

Antoine, L., Wolff, P., Westhof, E., Romby, P., & Marzi, S. (2019). Mapping post-transcriptional modifications in Staphylococcus aureus tRNAs by nanoLC/MSMS. *Biochimie, RNA Structure, Maturation*, Interactions and Functions, 164, 60–69. 10.1016/j.biochi.2019.07.003

Babosan, A., Fruchard, L., Krin, E., Carvalho, A., Mazel, D., & Baharoglu, Z. (2022). Nonessential tRNA and rRNA modifications impact the bacterial response to sub-MIC antibiotic stress. microLife, 3, uqac019. 10.1093/femsml/uqac019

Begik, O., Lucas, M. C., Pryszcz, L. P., Ramirez, J. M., Medina, R., Milenkovic, I., Cruciani, S., Liu, H., Vieira, H. G. S., Sas-Chen, A., Mattick, J. S., Schwartz, S., & Novoa, E. M. (2021). Quantitative profiling of pseudouridylation dynamics in native RNAs with nanopore sequencing. Nature Biotechnology, 39(10), Article 10. 10.1038/s41587-021-00915-6

Boccaletto, P., Stefaniak, F., Ray, A., Cappannini, A., Mukherjee, S., Purta, E., Kurkowska, M., Shirvanizadeh, N., Destefanis, E., Groza, P., Avşar, G., Romitelli, A., Pir, P., Dassi, E., Conticello, S. G., Aguilo, F., & Bujnicki, J. M. (2022). MODOMICS: A database of RNA modification pathways. 2021 update. Nucleic Acids Research, 50(D1), D231–D235. 10.1093/nar/gkab1083

Cruciani, S., Delgado-Tejedor, A., Pryszcz, L. P., Medina, R., Llovera, L., & Novoa, E. M. (2023). De novo basecalling of m6A modifications at single molecule and single nucleotide resolution (p. 2023.11.13.566801). bioRxiv. 10.1101/2023.11.13.566801

Cruciani, S., Delgado-Tejedor, A., Pryszcz, L. P., Medina, R., Llovera, L., & Novoa, E. M. (2025). De novo basecalling of RNA modifications at single molecule and nucleotide resolution. Genome Biology, 26(1), 38. 10.1186/s13059-025-03498-6

Cruciani, S., & Novoa, E. M. (2025). The new era of single-molecule RNA modification detection through nanopore base-calling models. Nature Reviews Molecular Cell Biology, 1–9. 10.1038/s41580-025-00896-3

Danecek, P., Bonfield, J. K., Liddle, J., Marshall, J., Ohan, V., Pollard, M. O., Whitwham, A., Keane, T., McCarthy, S. A., Davies, R. M., & Li, H. (2021). Twelve years of SAMtools and BCFtools. GigaScience, 10(2), giab008. 10.1093/gigascience/giab008

Delgado-Tejedor, A., Medina, R., Begik, O., Cozzuto, L., López, J., Blanco, S., Ponomarenko, J., & Novoa, E. M. (2024). Native RNA nanopore sequencing reveals antibiotic-induced loss of rRNA modifications in the A- and P-sites. Nature Communications, 15(1), 10054. 10.1038/s41467-024-54368-x

Diensthuber, G., Pryszcz, L., Llovera, L., Lucas, M. C., Delgado-Tejedor, A., Cruciani, S., Roignant, J.-Y., Begik, O., & Novoa, E. M. (2023). Enhanced detection of RNA modifications and mappability with high-accuracy nanopore RNA basecalling models (p. 2023.11.28.568965). bioRxiv. 10.1101/2023.11.28.568965

Dominissini, D., Nachtergaele, S., Moshitch-Moshkovitz, S., Peer, E., Kol, N., Ben-Haim, M. S., Dai, Q., Di Segni, A., Salmon-Divon, M., Clark, W. C., Zheng, G., Pan, T., Solomon, O., Eyal, E., Hershkovitz, V., Han, D., Doré, L. C., Amariglio, N., Rechavi, G., & He, C. (2016). The dynamic N1-methyladenosine methylome in eukaryotic messenger RNA. Nature, 530(7591), Article 7591. 10.1038/nature16998

Durant, P. C., Bajji, A. C., Sundaram, M., Kumar, R. K., & Davis, D. R. (2005). Structural Effects of Hypermodified Nucleosides in the Escherichia coli and Human tRNALys Anticodon Loop: The Effect of Nucleosides s2U, mcm5U, mcm5s2U, mnm5s2U, t6A, and ms2t6A. Biochemistry, 44(22), 8078–8089. 10.1021/bi050343f

Fleming, A. M., Bommisetti, P., Xiao, S., Bandarian, V., & Burrows, C. J. (2023a). Direct Nanopore Sequencing for the 17 RNA Modification Types in 36 Locations in the E. coli Ribosome Enables Monitoring of Stress-Dependent Changes. ACS Chemical Biology, 18(10), 2211–2223. 10.1021/acschembio.3c00166

Fleming, A. M., Bommisetti, P., Xiao, S., Bandarian, V., & Burrows, C. J. (2023b). Direct Nanopore Sequencing for the 17 RNA Modification Types in 36 Locations in the E. coli Ribosome Enables Monitoring of Stress-Dependent Changes. ACS Chemical Biology. 10.1021/acschembio.3c00166

Fonzino, A., Manzari, C., Spadavecchia, P., Munagala, U., Torrini, S., Conticello, S., Pesole, G., & Picardi, E. (2024). Unraveling C-to-U RNA editing events from direct RNA sequencing. RNA Biology, 21(1), 1–14. 10.1080/15476286.2023.2290843

Gamaarachchi, H., Lam, C. W., Jayatilaka, G., Samarakoon, H., Simpson, J. T., Smith, M. A., & Parameswaran, S. (2020). GPU accelerated adaptive banded event alignment for rapid comparative nanopore signal analysis. BMC Bioinformatics, 21(1), 343. 10.1186/s12859-020-03697-x

Gao, Y., Liu, X., Wu, B., Wang, H., Xi, F., Kohnen, M. V., Reddy, A. S. N., & Gu, L. (2021). Quantitative profiling of N6-methyladenosine at single-base resolution in stem-differentiating xylem of Populus trichocarpa using Nanopore direct RNA sequencing. Genome Biology, 22(1), 22. 10.1186/s13059-020-02241-7

Grozhik, A. V., Linder, B., Olarerin-George, A. O., & Jaffrey, S. R. (2017). Mapping m6A at Individual-Nucleotide Resolution Using Crosslinking and Immunoprecipitation (miCLIP). In A. Lusser (Ed.), RNA Methylation: Methods and Protocols (pp. 55–78). Springer. 10.1007/978-1-4939-6807-7_5

Guo, Z., Shao, Y., Tan, L., Lu, B., Deng, X., Chen, S., & Li, R. (2025). Enhanced detection of RNA modifications in Escherichia coli utilizing direct RNA sequencing. Cell Reports Methods, 5(9). 10.1016/j.crmeth.2025.101168

Hassan, D., Acevedo, D., Daulatabad, S. V., Mir, Q., & Janga, S. C. (2021). Penguin: A Tool for Predicting Pseudouridine Sites in Direct RNA Nanopore Sequencing Data. bioRxiv, 2021.03.31.437901. 10.1101/2021.03.31.437901

Hoernes, T. P., Clementi, N., Faserl, K., Glasner, H., Breuker, K., Lindner, H., Hüttenhofer, A., & Erlacher, M. D. (2016). Nucleotide modifications within bacterial messenger RNAs regulate their translation and are able to rewire the genetic code. Nucleic Acids Research, 44(2), 852–862. 10.1093/nar/gkv1182

Jain, M., Abu-Shumays, R., Olsen, H. E., & Akeson, M. (2022). Advances in nanopore direct RNA sequencing. Nature Methods, 19(10), Article 10. 10.1038/s41592-022-01633-w

Jenjaroenpun, P., Wongsurawat, T., Wadley, T. D., Wassenaar, T. M., Liu, J., Dai, Q., Wanchai, V., Akel, N. S., Jamshidi-Parsian, A., Franco, A. T., Boysen, G., Jennings, M. L., Ussery, D. W., He, C., & Nookaew, I. (2021). Decoding the epitranscriptional landscape from native RNA sequences. Nucleic Acids Research, 49(2), e7. 10.1093/nar/gkaa620

Leger, A., Amaral, P. P., Pandolfini, L., Capitanchik, C., Capraro, F., Miano, V., Migliori, V., Toolan-Kerr, P., Sideri, T., Enright, A. J., Tzelepis, K., van Werven, F. J., Luscombe, N. M., Barbieri, I., Ule, J., Fitzgerald, T., Birney, E., Leonardi, T., & Kouzarides, T. (2021). RNA modifications detection by comparative Nanopore direct RNA sequencing. Nature Communications, 12(1), Article 1. 10.1038/s41467-021-27393-3

Li, H. (2018). Minimap2: Pairwise alignment for nucleotide sequences. Bioinformatics, 34(18), 3094–3100. 10.1093/bioinformatics/bty191

Li, H., Handsaker, B., Wysoker, A., Fennell, T., Ruan, J., Homer, N., Marth, G., Abecasis, G., & Durbin, R. (2009). The Sequence Alignment/Map format and SAMtools. Bioinformatics, 25(16), 2078–2079. 10.1093/bioinformatics/btp352

Li, X., Zhu, P., Ma, S., Song, J., Bai, J., Sun, F., & Yi, C. (2015). Chemical pulldown reveals dynamic pseudouridylation of the mammalian transcriptome. Nature Chemical Biology, 11(8), Article 8. 10.1038/nchembio.1836

Liu, H., Begik, O., Lucas, M. C., Ramirez, J. M., Mason, C. E., Wiener, D., Schwartz, S., Mattick, J. S., Smith, M. A., & Novoa, E. M. (2019). Accurate detection of m6A RNA modifications in native RNA sequences. Nature Communications, 10(1), Article 1. 10.1038/s41467-019-11713-9

Lorenz, D. A., Sathe, S., Einstein, J. M., & Yeo, G. W. (2020). Direct RNA sequencing enables m6A detection in endogenous transcript isoforms at base-specific resolution. RNA, 26(1), 19–28. 10.1261/rna.072785.119

Lucas, M. C., Pryszcz, L. P., Medina, R., Milenkovic, I., Camacho, N., Marchand, V., Motorin, Y., Ribas de Pouplana, L., & Novoa, E. M. (2024). Quantitative analysis of tRNA abundance and modifications by nanopore RNA sequencing. Nature Biotechnology, 42(1), Article 1. 10.1038/s41587-023-01743-6

Luo, T., Xu, M., Wang, M., Chen, F., & Shi, J. (2026). Systematic evaluation of computational tools for multitype RNA modification detection using nanopore direct RNA sequencing. Nature Methods, 23(2), 438–450. 10.1038/s41592-025-02974-y

Maestri, S., Furlan, M., Mulroney, L., Coscujuela Tarrero, L., Ugolini, C., Dalla Pozza, F., Leonardi, T., Birney, E., Nicassio, F., & Pelizzola, M. (2024). Benchmarking of computational methods for m6A profiling with Nanopore direct RNA sequencing. Briefings in Bioinformatics, 25(2), bbae001. 10.1093/bib/bbae001

Maier, K. C., Gressel, S., Cramer, P., & Schwalb, B. (2020). Native molecule sequencing by nano-ID reveals synthesis and stability of RNA isoforms. Genome Research, 30(9), 1332–1344. 10.1101/gr.257857.119

Marchand, V., Ayadi, L., Ernst, F. G. M., Hertler, J., Bourguignon-Igel, V., Galvanin, A., Kotter, A., Helm, M., Lafontaine, D. L. J., & Motorin, Y. (2018). AlkAniline-Seq: Profiling of m7G and m3C RNA Modifications at Single Nucleotide Resolution. Angewandte Chemie International Edition, 57(51), 16785–16790. 10.1002/anie.201810946

Martinek, V., Martin, J., Belair, C., Payea, M. J., Malla, S., Alexiou, P., & Maragkakis, M. (2024). Deep learning and direct sequencing of labeled RNA captures transcriptome dynamics. NAR Genomics and Bioinformatics, 6(3), lqae116. 10.1093/nargab/lqae116

Mateos, P. A., Sethi, A. J., Ravindran, A., Srivastava, A., Kanchi, M., Mahmud, S., Guarnacci, M., Woodward, K., Xu, J., Yuen, Z. W. S., Zhou, Y., Sneddon, A., Hamilton, W., Gao, J., Starrs, L. M., Hayashi, R., Wickramasinghe, V., Zarnack, K., Preiss, T., … Eyras, E. (2024). Simultaneous identification of m6A and m5C reveals coordinated RNA modification at single-molecule resolution (p. 2022.03.14.484124). bioRxiv. 10.1101/2022.03.14.484124

Morampalli, B. R., Silander, O., Morampalli, B. R., & Silander, O. (2021). Synthesis of in vitro transcribed RNA from whole bacterial transcriptome. https://www.protocols.io/view/synthesis-of-in-vitro-transcribed-rna-from-whole-b-brbmm2k6

Naarmann-de Vries, I. S., Zorbas, C., Lemsara, A., Piechotta, M., Ernst, F. G. M., Wacheul, L., Lafontaine, D. L. J., & Dieterich, C. (2023). Comprehensive identification of diverse ribosomal RNA modifications by targeted nanopore direct RNA sequencing and JACUSA2. RNA Biology, 20(1), 652–665. 10.1080/15476286.2023.2248752

Nguyen, T. A., Heng, J. W. J., Kaewsapsak, P., Kok, E. P. L., Stanojević, D., Liu, H., Cardilla, A., Praditya, A., Yi, Z., Lin, M., Aw, J. G. A., Ho, Y. Y., Peh, K. L. E., Wang, Y., Zhong, Q., Heraud-Farlow, J., Xue, S., Reversade, B., Walkley, C., … Tan, M. H. (2022a). Direct identification of A-to-I editing sites with nanopore native RNA sequencing. Nature Methods, 19(7), Article 7. 10.1038/s41592-022-01513-3

Nguyen, T. A., Heng, J. W. J., Kaewsapsak, P., Kok, E. P. L., Stanojević, D., Liu, H., Cardilla, A., Praditya, A., Yi, Z., Lin, M., Aw, J. G. A., Ho, Y. Y., Peh, K. L. E., Wang, Y., Zhong, Q., Heraud-Farlow, J., Xue, S., Reversade, B., Walkley, C., … Tan, M. H. (2022b). Direct identification of A-to-I editing sites with nanopore native RNA sequencing. Nature Methods, 19(7), Article 7. 10.1038/s41592-022-01513-3

Ohnezeit, D., Loliashvili, E., Putzel, G., Verstraten, R., Liu, J., Nicholson, L. S., Pironti, A., Jaffrey, S. R., Depledge, D. P., & Wilson, A. C. (2026). Calibrated analysis framework for nanopore direct RNA sequencing uncovers cell-specific m6A stoichiometry at conserved sites (p. 2025.11.02.686099). bioRxiv. 10.1101/2025.11.02.686099

Parker, M. T., Barton, G. J., & Simpson, G. G. (2021a). Yanocomp: Robust prediction of m6A modifications in individual nanopore direct RNA reads (p. 2021.06.15.448494). 10.1101/2021.06.15.448494

Parker, M. T., Barton, G. J., & Simpson, G. G. (2021b). Yanocomp: Robust prediction of m6A modifications in individual nanopore direct RNA reads (p. 2021.06.15.448494). bioRxiv. 10.1101/2021.06.15.448494

Parker, M. T., Knop, K., Sherwood, A. V., Schurch, N. J., Mackinnon, K., Gould, P. D., Hall, A. J., Barton, G. J., & Simpson, G. G. (2020). Nanopore direct RNA sequencing maps the complexity of Arabidopsis mRNA processing and m6A modification. eLife, 9, e49658. 10.7554/eLife.49658

Patiño-Guillén, G., Pešović, J., Panić, M., Savić-Pavićević, D., Bošković, F., & Keyser, U. F. (2024). Single-molecule RNA sizing enables quantitative analysis of alternative transcription termination. Nature Communications, 15(1), Article 1. 10.1038/s41467-024-45968-8

Piechotta, M., Naarmann-de Vries, I. S., Wang, Q., Altmüller, J., & Dieterich, C. (2022). RNA modification mapping with JACUSA2. Genome Biology, 23(1), 115. 10.1186/s13059-022-02676-0

Popova, A. M., & Williamson, J. R. (2014). Quantitative Analysis of rRNA Modifications Using Stable Isotope Labeling and Mass Spectrometry. Journal of the American Chemical Society, 136(5), 2058–2069. 10.1021/ja412084b

Pratanwanich, P. N., Yao, F., Chen, Y., Koh, C. W. Q., Wan, Y. K., Hendra, C., Poon, P., Goh, Y. T., Yap, P. M. L., Chooi, J. Y., Chng, W. J., Ng, S. B., Thiery, A., Goh, W. S. S., & Göke, J. (2021). Identification of differential RNA modifications from nanopore direct RNA sequencing with xPore. Nature Biotechnology, 39(11), Article 11. 10.1038/s41587-021-00949-w

Pust, M.-M., Davenport, C. F., Wiehlmann, L., & Tümmler, B. (2022). Direct RNA Nanopore Sequencing of Pseudomonas aeruginosa Clone C Transcriptomes. Journal of Bacteriology, 204(1), e00418–21. 10.1128/JB.00418-21

Qin, H., Ou, L., Gao, J., Chen, L., Wang, J.-W., Hao, P., & Li, X. (2022). DENA: Training an authentic neural network model using Nanopore sequencing data of Arabidopsis transcripts for detection and quantification of N6-methyladenosine on RNA. Genome Biology, 23(1), 25. 10.1186/s13059-021-02598-3

Riquelme-Barrios, S., Vásquez-Camus, L., Cusack, S. A., Burdack, K., Petrov, D. P., Yeşiltaç-Tosun, G. N., Kaiser, S., Giehr, P., & Jung, K. (2025). Direct RNA sequencing of the Escherichia coli epitranscriptome uncovers alterations under heat stress. Nucleic Acids Research, 53(6), gkaf175. 10.1093/nar/gkaf175

Schaefer, M., Pollex, T., Hanna, K., & Lyko, F. (2009). RNA cytosine methylation analysis by bisulfite sequencing. Nucleic Acids Research, 37(2), e12. 10.1093/nar/gkn954

Shen, W., Le, S., Li, Y., & Hu, F. (2016). SeqKit: A Cross-Platform and Ultrafast Toolkit for FASTA/Q File Manipulation. PLOS ONE, 11(10), e0163962. 10.1371/journal.pone.0163962

Smith, M. A., Ersavas, T., Ferguson, J. M., Liu, H., Lucas, M. C., Begik, O., Bojarski, L., Barton, K., & Novoa, E. M. (2020). Molecular barcoding of native RNAs using nanopore sequencing and deep learning. Genome Research, 30(9), 1345–1353. 10.1101/gr.260836.120

Sordyl, D., Boileau, E., Bernat, A., Maiti, S., Mukherjee, S., Moafinejad, S. N., Farsani, M. A., Shavina, A., Cappannini, A., Agostini, G., Conticello, S. G., Stefaniak, F., Dieterich, C., Purta, E., & Bujnicki, J. M. (2026). MODOMICS: A database of RNA modifications and related information. 2025 update and 20th anniversary. Nucleic Acids Research, 54(D1), D219–D225. 10.1093/nar/gkaf1284

Spangenberg, J., Mündnich, S., Busch, A., Pastore, S., Wierczeiko, A., Goettsch, W., Dietrich, V., Pryszcz, L. P., Cruciani, S., Novoa, E. M., Joshi, K., Perera, R., Di Giorgio, S., Arrubarrena, P., Tellioglu, I., Poon, C.-L., Wan, Y. K., Göke, J., Hildebrandt, A., … Alagna, N. (2025). The RMaP challenge of predicting RNA modifications by nanopore sequencing. Communications Chemistry, 8(1), 115. 10.1038/s42004-025-01507-0

Stoiber, M., Quick, J., Egan, R., Lee, J. E., Celniker, S., Neely, R. K., Loman, N., Pennacchio, L. A., & Brown, J. (2017). De novo Identification of DNA Modifications Enabled by Genome-Guided Nanopore Signal Processing (p. 094672). bioRxiv. 10.1101/094672

Tan, L., Guo, Z., Shao, Y., Ye, L., Wang, M., Deng, X., Chen, S., & Li, R. (2024). Analysis of bacterial transcriptome and epitranscriptome using nanopore direct RNA sequencing. Nucleic Acids Research, gkae601. 10.1093/nar/gkae601

Tan, L., Guo, Z., Wang, X., Kim, D. Y., & Li, R. (2024a). Utilization of nanopore direct RNA sequencing to analyze viral RNA modifications. mSystems, 0(0), e01163–23. 10.1128/msystems.01163-23

Tan, L., Guo, Z., Wang, X., Kim, D. Y., & Li, R. (2024b). Utilization of nanopore direct RNA sequencing to analyze viral RNA modifications. mSystems, 9(2), e01163–23. 10.1128/msystems.01163-23

Teng, H., Stoiber, M., Bar-Joseph, Z., & Kingsford, C. (2024). Detecting m6A RNA modification from nanopore sequencing using a semi-supervised learning framework (p. 2024.01.06.574484). bioRxiv. 10.1101/2024.01.06.574484

Ueda, H. (2021). nanoDoc: RNA modification detection using Nanopore raw reads with Deep One-Class Classification (p. 2020.09.13.295089). bioRxiv. 10.1101/2020.09.13.295089

White, L. K., Dobson, K., Pozo, S. del, Bilodeaux, J. M., Andersen, S. E., Baldwin, A., Barrington, C., Körtel, N., Martinez-Seidel, F., Strugar, S. M., Watt, K. E. N., Mukherjee, N., & Hesselberth, J. R. (2024). Comparative analysis of 43 distinct RNA modifications by nanopore tRNA sequencing (p. 2024.07.23.604651). bioRxiv. 10.1101/2024.07.23.604651

White, L. K., & Hesselberth, J. R. (2022). Modification mapping by nanopore sequencing. Frontiers in Genetics, 13. https://www.frontiersin.org/articles/10.3389/fgene.2022.1037134

Wick, R. (2022). Rrwick/Filtlong [C++]. https://github.com/rrwick/Filtlong (Original work published 2017)

Workman, R. E., Tang, A. D., Tang, P. S., Jain, M., Tyson, J. R., Razaghi, R., Zuzarte, P. C., Gilpatrick, T., Payne, A., Quick, J., Sadowski, N., Holmes, N., de Jesus, J. G., Jones, K. L., Soulette, C. M., Snutch, T. P., Loman, N., Paten, B., Loose, M., … Timp, W. (2019). Nanopore native RNA sequencing of a human poly(A) transcriptome. Nature Methods, 16(12), Article 12. 10.1038/s41592-019-0617-2

Wu, Y., Shao, W., Yan, M., Wang, Y., Xu, P., Huang, G., Li, X., Gregory, B. D., Yang, J., Wang, H., & Yu, X. (2024). Transfer learning enables identification of multiple types of RNA modifications using nanopore direct RNA sequencing. Nature Communications, 15(1), 4049. 10.1038/s41467-024-48437-4

Xuan, J., Chen, L., Chen, Z., Pang, J., Huang, J., Lin, J., Zheng, L., Li, B., Qu, L., & Yang, J. (2024). RMBase v3.0: Decode the landscape, mechanisms and functions of RNA modifications. Nucleic Acids Research, 52(D1), D273–D284. 10.1093/nar/gkad1070

Zhao, X., Zhang, Y., Hang, D., Meng, J., & Wei, Z. (2022). Detecting RNA modification using direct RNA sequencing: A systematic review. Computational and Structural Biotechnology Journal, 20, 5740–5749. 10.1016/j.csbj.2022.10.023

Zhong, Z.-D., Xie, Y.-Y., Chen, H.-X., Lan, Y.-L., Liu, X.-H., Ji, J.-Y., Wu, F., Jin, L., Chen, J., Mak, D. W., Zhang, Z., & Luo, G.-Z. (2023a). Systematic comparison of tools used for m6A mapping from nanopore direct RNA sequencing. Nature Communications, 14(1), Article 1. 10.1038/s41467-023-37596-5

Zhong, Z.-D., Xie, Y.-Y., Chen, H.-X., Lan, Y.-L., Liu, X.-H., Ji, J.-Y., Wu, F., Jin, L., Chen, J., Mak, D. W., Zhang, Z., & Luo, G.-Z. (2023b). Systematic comparison of tools used for m6A mapping from nanopore direct RNA sequencing. Nature Communications, 14(1), 1906. 10.1038/s41467-023-37596-5

Zou, Y., Ahsan, M. U., Chan, J., Meng, W., Gao, S.-J., Huang, Y., & Wang, K. (2025). A comparative evaluation of computational models for RNA modification detection using nanopore sequencing with RNA004 chemistry. Briefings in Bioinformatics, 26(4), bbaf404. 10.1093/bib/bbaf404

